# Macrophage-Dendritic Cell-T-Cell Tetrads Orchestrate Antitumor Immunity and Response to Checkpoint Blockade

**DOI:** 10.64898/2025.12.24.696419

**Authors:** Mehdi Chaib, Muhammad Aminu, Shelley Herbrich, Mahshid Arabi, Yue Xuan, Akshay Basi, Anna Casasent, Marc D. Macaluso, Kenneth H. Hu, Matthew Gubin, Xi Chen, James J. Mancuso, Sreyashi Basu, Sonali Jindal, Jared Burks, Stephanie Watowich, Jia Wu, James P. Allison, Padmanee Sharma

**Author notes:** These authors contributed equally (Mehdi Chaib, Muhammad Aminu).

## Abstract

Immune checkpoint inhibitors (ICIs) elicit durable responses in only a subset of patients with solid tumors, underscoring the need to define the cellular architectures that govern effective antitumor immunity. Here we identify a spatially organized multicellular immune unit comprising macrophages, cDC1s, CD4⁺ T-cells, and CD8⁺ T-cells that emerges in response to anti-CTLA-4 or dual checkpoint blockade. We term these structures *tetrads*. Using multiplexed imaging and spatial transcriptomics in mouse and human tumors, we show that tetrads assemble early during immune priming, depend on the ICOS–ICOSL pathway, and are enriched for ICOS⁺ Th1-like CD4⁺ T cells and ICOSL^high^ cDC1s. CD8⁺ T-cells within tetrads exhibit an activated, non-terminally differentiated state, while tetrad-associated macrophages display an interferon-γ–responsive program that sustains CD8⁺ T-cell function and prevents dysfunction. Functionally, ICOSL⁺ cDC1s are required for tumor eradication *in vivo*. In patients with bladder cancer treated with neoadjuvant dual checkpoint blockade, tetrad, but not triad or dyad formation correlates with clinical response. These findings establish tetrads as a fundamental cellular unit coordinating antitumor immunity and responsiveness to ICIs.

## Main

Immunotherapy, including immune checkpoint inhibitors (ICI), has transformed the landscape of cancer treatment; however, only a subset of patients with solid tumors responds to these therapies. Interactions among neighboring cells within the tumor microenvironment (TME) critically determine the outcome of immunotherapy ^1^. Thus, multicellular networks organized in space within solid tumors are key orchestrators of effective antitumor immunity. Most clinically applied cancer immunotherapies rely on the ability of cytotoxic CD8⁺ T cells to directly engage tumor cells, yet a growing body of evidence underscores the critical role of CD4⁺ T cells in generating durable antitumor immune responses ^2^. CD4⁺ T cells can target tumor cells directly through cytolytic mechanisms or indirectly by modulating the TME ^3^. One major indirect pathway is the cDC1-CD4⁺ T cell axis, in which cDC1s both prime and are “licensed” by CD4⁺ T cells through CD40-CD40L interactions to elicit effective antitumor immunity ^4^.

Recent studies have advanced our understanding of the spatial and functional coordination between dendritic cells (DCs) and T cells. In mice, optimal antitumor immunity is enhanced when epitopes recognized by CD4⁺ and CD8⁺ T cells be linked and presented by the same antigen-presenting cell (APC) ^5^. This led to the emergence of the three-cell-type hypothesis or “triads” ^6^. More recent studies showed that these triads are required for effective tumor eradication and the presence of CD4 T cells within triads reprograms CD8 T cells and prevents their terminal differentiation ^7^. In addition, triads are associated with a better prognosis in cancer patients and predict response to ICI ^7,8^. Yet, the precise cellular subtypes that form these structures, the pathways governing their assembly and function, and whether they are actively induced by ICI remain unresolved.

We and others have shown that anti-CTLA-4 and anti-PD-1 therapies engage distinct yet complementary cellular mechanisms ^9–11^. Whereas anti-PD-1 primarily reinvigorates T cells during later stages of activation, anti-CTLA-4 acts earlier by restoring costimulation via APCs during the priming phase. Notably, anti-CTLA-4, but not anti-PD-1, selectively expands inducible T cell costimulatory (ICOS)⁺ Th1-like CD4⁺ effector cells in mice and humans ^9,12–15^. We and others have also shown that the ICOS-ICOS ligand (ICOSL) pathway plays a crucial role in the antitumor effect of anti-CTLA-4 therapy ^16^, which requires macrophages ^17^. These complementary mechanisms explain the enhanced clinical efficacy of dual checkpoint blockade, as anti-CTLA-4 promotes T cell priming while anti-PD-1 sustains T cell effector function, thereby broadening the window of immune activation. This synergy is reflected in the superior survival benefit observed in melanoma patients receiving combination therapy compared to monotherapy ^18,19^. Understanding the mechanisms by which anti-CTLA-4 shapes early immune priming is therefore critical for defining the quality and diversity of the ensuing T cell response.

Although macrophages are often linked to poor prognosis and therapeutic resistance ^20^, emerging evidence indicates that specific macrophage subsets are important to mount an effective antitumor immune response ^21,22^. Tumor-associated macrophages can adopt either a wound-healing, anti-inflammatory phenotype that promotes tumor progression and therapy resistance, or a pro-inflammatory, immunostimulatory phenotype that supports antitumor immune responses ^20,23^. However, how macrophages, DCs, and T cells interact spatially within tumors in response to ICI remains poorly understood. Using multiplexed imaging and spatial transcriptomics, we identified the formation of macrophage-cDC1-CD4-CD8 T cell clusters in response to anti-CTLA-4 or dual checkpoint blockade in mouse and human tumors. We termed these interacting cell clusters “tetrads”. Critically, most cDC1-CD4-CD8 T-cell intra-tumoral triads engaged neighboring macrophages following anti-CTLA-4 treatment, suggesting that “tetrads,” rather than triads, assemble early during the intratumoral priming phase in response to anti-CTLA-4. CD8⁺ T cells within tetrads display an activated phenotype distinct from terminally differentiated counterparts outside of tetrads. Mechanistically, tetrad formation depends on the ICOS-ICOSL pathway, marked by enrichment of ICOS⁺ Th1-like CD4⁺ T cells and ICOSL^high^ cDC1s within tetrad regions. Functionally, adoptive transfer of ICOSL⁺ cDC1s in combination with anti-CTLA-4 therapy eradicated ∼50% of B16F10 tumors, whereas ICOSL⁻^/^⁻ cDC1s failed to mediate tumor clearance. Interestingly, tetrad-associated macrophages exhibit an interferon-γ-responsive transcriptional program and sustain CD8⁺ T cell functionality by preventing terminal differentiation. Strikingly, tetrad formation, but not triad or dyad formations in tumors from bladder cancer patients treated with neoadjuvant dual checkpoint blockade correlated with clinical response and mirrored mechanisms observed in mice. These findings identify tetrads as a fundamental cellular unit that orchestrates intratumoral T cell priming, tumor elimination, and responsiveness to ICI.

### Identification of immune tetrads induced by anti-CTLA-4 and dual checkpoint inhibition therapies in murine tumors

To visualize APC-T cell interactions in tumors, we used multiplexed imaging platforms (Lunaphore COMET and RNAscope), which enable simultaneous visualization of up to 20 protein and 12 RNA targets within a single formalin-fixed paraffin-embedded (FFPE) tissue section at subcellular resolution. B16F10 tumors were collected on day 12 after tumor implantation from C57BL/6J mice treated with two doses on days 7 and 10 of anti-CTLA-4, anti-PD-1, combination therapy, or left untreated. We generated tissue microarrays (TMAs) by randomly selecting two 1 mm-diameter regions of interest (ROIs) from each tumor, yielding a total of 108 tumor cores across treatment groups. Multiplex panels included up to 20 protein markers capturing major and minor immune cell populations (CD3, CD45, CD4, CD8, CD103, CD11c, F4/80, TOX, TCF1, etc.) as well as up to 12 RNA probes to detect genes of secreted factors and transcription factors such as *Cxcl9* and *Isg15,* respectively. The full list of protein and RNA targets is listed in the methods section. Raw images were processed and analyzed using a validated, data-driven computational pipeline that performs unsupervised pixel clustering to define major cell types with high resolution ^24^. We then developed custom algorithms that applied cell-mask segmentation to delineate cell boundaries and identify cell-cell interactions, classifying cells as interacting only when in direct membrane contact (Fig. 1A).

**Figure 1:**
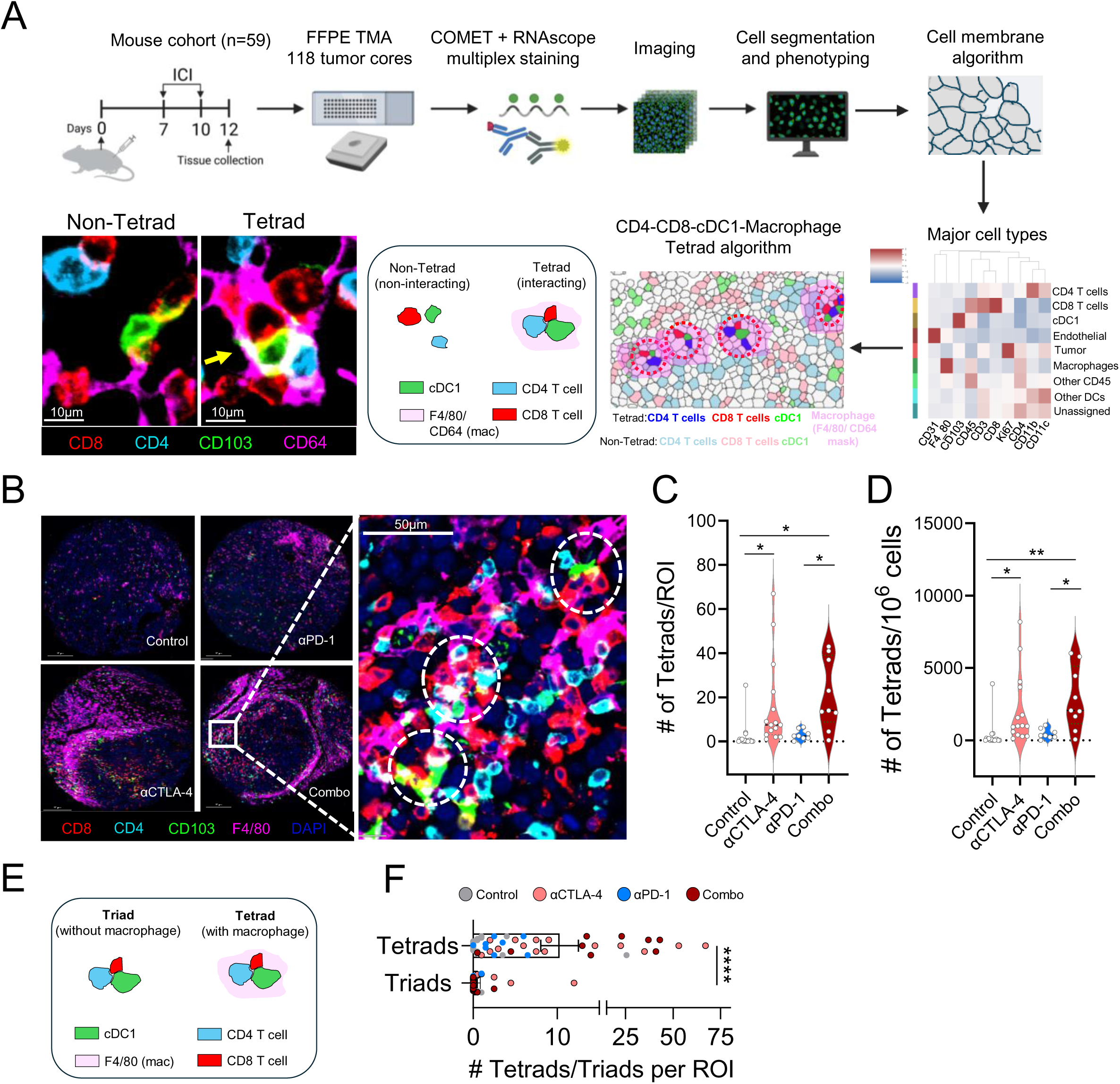
Identification of immune tetrads induced by anti-CTLA-4 and dual checkpoint inhibition therapies in murine tumors. **(A)** Schematics illustrating the sequential immunofluorescence and RNAscope pipeline analysis for major cell phenotyping and tetrad identification in mouse B16F10 tumors collected on day 12 (n = 59 total biological replicates for entire study for B16F10 tumor models). **(B)** Representative COMET images highlighting tetrads (white circles). Tetrads are composed of CD8^+^ T-cells (red), CD4^+^ T-cells (blue cyan), cDC1 (green) and macrophages (purple). Nuclear staining (DAPI) is highlighted in dark blue. **(C-D)** Tetrad quantification representing absolute numbers per ROI **(C)**, and numbers normalized to the total number of cells per each ROI **(D)**. Data are pooled from 3 independent experiments (n = 8-16 biological replicates per group). Significance is calculated by One-way ANOVA with Tukey’s multiple comparisons test. (\**p*<0.05, \*\**p*<0.01). Values are mean ± SEM. **(E-F)** Triad and tetrad comparison in mouse B16F10 tumors. **(E)** Schematics illustrating differences between triad and tetrad quantification (mutually exclusive). **(F)** Number of triads and tetrads per ROI (color-coded per treatment group). Data are pooled from 3 independent experiments (n = 48 biological replicates). (****p<0.0001, unpaired T-test). Values are mean ± SEM.

Unexpectedly, we observed that canonical cDC1-CD4-CD8 T-cell triads frequently incorporated a fourth interacting cell type—macrophages. We termed these four-cell structures “tetrads.” To systematically quantify tetrads, we designed an algorithm requiring that cDC1s, CD4⁺ T cells, and CD8⁺ T cells be directly interacting and that all three cell types share contact with a macrophage mask (F4/80⁺ or CD64⁺), indicating engagement with a neighboring macrophage. Tetrad formation was markedly increased in tumors treated with anti-CTLA-4 or dual checkpoint blockade (anti-CTLA-4 + anti-PD-1) compared to anti-PD-1 monotherapy or untreated controls (Fig. 1B-D), both in absolute numbers per ROI (Fig. 1C) and when normalized to total cell counts (Fig. 1D). Notably, when comparing triad versus tetrad interactions, the majority of cDC1-CD4-CD8 T-cell triads included macrophages, indicating that four-cell “tetrads,” rather than classical triads, dominate in response to anti-CTLA-4-based therapy (Fig. 1E,F). These findings suggest that anti-CTLA-4 and dual checkpoint inhibition promote coordinated interactions among macrophages, cDC1s, CD4⁺ T cells, and CD8⁺ T cells, forming a distinct multicellular structure—the tetrad—that differs from previously described triads.

Because B16F10 tumors are only partially responsive to ICI, exhibiting delayed but incomplete tumor regression ^17,25^, we next asked whether ICI-sensitive tumors display higher tetrad abundance. We analyzed the Y1.7LI (mLama) melanoma model, derived from Braf^V600E^ Pten^−/−^ Cdkn2a^−/−^ YUMM1.7 cells engineered to express defined MHC-I (mLama4) and MHC-II (mItgb1) neoantigens, rendering it ICI-sensitive where most tumors are rejected post-treatment ^26^. Strikingly, mLama tumors exhibited substantially higher baseline tetrad density compared to either control or anti-CTLA-4-treated B16F10 tumors (Fig. S1A-C). These results indicate that while anti-CTLA-4 therapy can induce tetrad formation, pre-existing tetrads may potentiate responsiveness to ICI. Collectively, these findings support a four-cell-type hypothesis, in which macrophage-cDC1-CD4-CD8 T-cell tetrads emerge early during the priming phase and serve as predictors of therapeutic response to immune checkpoint inhibition.

### Tetrad-associated CD8⁺ T cells display features of activation and effector differentiation

Dysfunctional or terminally differentiated tumor-infiltrating CD8⁺ T cells, oten referred to as exhausted, typically express high levels of inhibitory receptors such as PD-1, TIM-3, and LAG-3, along with the transcription factor TOX, while exhibiting diminished production of effector molecules including IFNγ, TNFα, and granzyme B (GZMB)--hallmarks of activated effector cytotoxic T cells. In contrast, stem-like progenitor CD8⁺ T cells expressing the transcription factor TCF1 serve as a reservoir for effector differentiation, and the presence of both populations correlates with improved prognosis in cancer patients ^27^. To define the phenotype of tetrad-associated CD8⁺ T cells, we analyzed their spatial distribution and transcriptional state relative to tetrads. Specifically, we defined 50-μm and 100-μm radii surrounding tetrads to assess CD8⁺ T-cell phenotypes within, adjacent to, and distal from these structures.

We subclustered all CD8⁺ T cells based on transcription factor and cell division marker expression (proteins - TOX, Ki67, and TCF1) and functional markers (RNA - *Tnfa, Gzmb*, and *Ifng*) (Fig. 2A). Six distinct CD8⁺ T-cell clusters were identified: clusters C0 and C4 expressed high TOX, consistent with a terminally differentiated or dysfunctional phenotype; cluster C2 displayed an effector-activated signature characterized by elevated *Gzmb* and *Ifng* expression; and cluster C5 represented a stem-like subset with high TCF1 expression. Clusters C1 and C3 exhibited mixed or intermediate features and were classified as “other” (Fig. 2B,C). We observed a significantly higher frequency of the *Ifng*^hi^ *Gzmb*^hi^ effector-like cluster (C2) within tetrads compared to regions outside tetrads, whereas the TOX^hi^ terminally differentiated-like cluster (C4) was markedly enriched outside tetrads (Fig. 2D). The stem-like TCF1^hi^ cluster (C5) was also significantly enriched within tetrads, while cluster C1 predominated outside these structures (Fig. 2D). Spatial gradient analysis revealed that only clusters C2 and C4 exhibited a distance-dependent distribution: effector-like C2 CD8^+^ T cells were most abundant within tetrads, intermediate at 50 μm and 100 μm from tetrads, and lowest beyond 100 μm (Fig. 2E). In contrast, TOX^hi^ cluster C4 cells were lower within tetrads and progressively enriched with increasing distance (Fig. 2F). Other clusters did not display similar diffusion patterns (Fig. S2A-D). These results indicate that effector-like and stem-like CD8⁺ T cells are preferentially localized within tetrads, whereas terminally differentiated, dysfunctional CD8⁺ T cells accumulate outside these immune assemblies.

**Figure 2:**
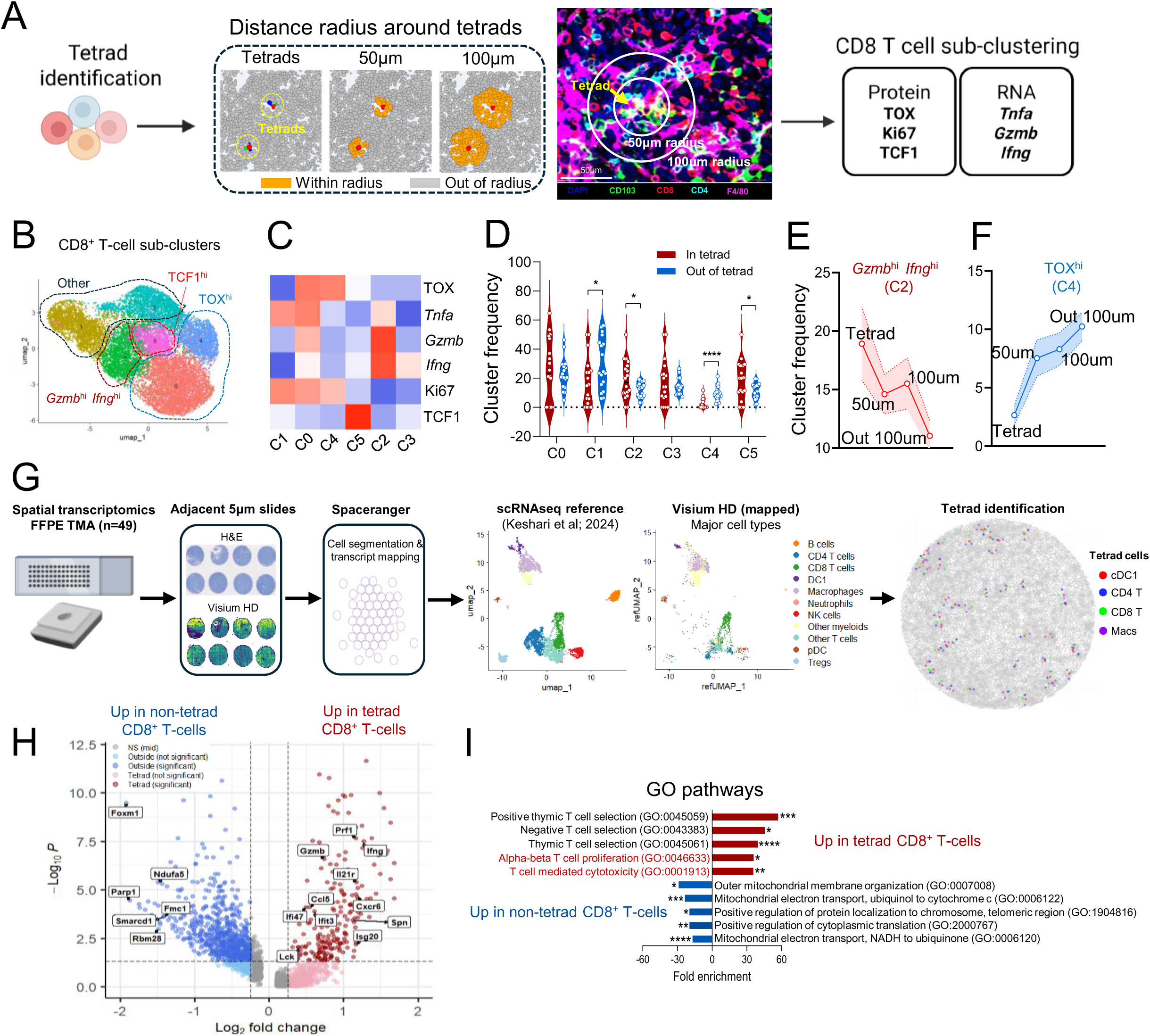
Tetrad-associated CD8⁺ T cells display features of activation and effector differentiation. **(A)** Schematic illustrating the analytical workflow for sub-clustering CD8⁺ T-cells associated with tetrads and within 50-μm and 100-μm neighborhoods surrounding tetrads in mouse B16F10 tumors. **(B-C)** UMAP **(B)** and heatmap **(C)** highlighting different CD8^+^ T-cell sub-clusters and marker expression from panel A. **(D)** CD8^+^ T-cell sub-clusters frequency in tetrad versus outside of tetrads. (n = 18 biological replicates). (\**p*<0.05, \*\*\*\**p*<0.0001, unpaired T-test). Values are mean ± SEM. **(E-F)** C2 (*Gzmb*^hi^ *Ifng*^hi^) **(E)** and C4 (TOX^hi^) **(F)** CD8^+^ T-cell sub-cluster frequency distribution in tetrads, within 50-μm and 100-μm neighborhoods, and outside of 100-μm. (n = 18 biological replicates) **(G)** Schematic illustrating Visium HD data analysis pipeline of mouse B16F10 tumors (n = 48 biological replicates). **(H)** Volcano plot for DEGs between tetrad (red) and non-tetrad (blue) CD8^+^ T-cells. Statistically significantly DEGs (*p* < 0.05) are shown in red and blue, with select genes highlighted for reference. **(I)** Top 5 GO terms enriched for genes upregulated in tetrad (red) and non-tetrad (blue) CD8^+^ T-cells.

To further characterize the molecular features of tetrad-associated CD8⁺ T cells at higher resolution, we performed spatial transcriptomic profiling using Visium HD on *n* = 49 B16F10 tumor cores. Visium HD captures whole-transcriptome data with near-single-cell resolution. We followed an established pipeline for tissue data processing and analysis, using a 5-μm adjacent section for H&E-based cell segmentation^28^. To define major immune populations, we reanalyzed a publicly available single-cell RNA-seq dataset ^29^, and transferred the expression profiles of the 2,000 most representative genes per immune cell type into the Visium HD dataset to spatially map immune subsets within tumors (Fig. 2G).

Following tetrad identification, we compared the transcriptomic profiles of CD8⁺ T cells located within tetrad regions versus those outside tetrads. Differentially expressed genes (DEGs) upregulated in tetrad-associated CD8⁺ T cells included canonical cytotoxic effector genes such as *Prf1*, *Gzmb*, and *Ifng* (Fig. 2H), consistent with our multiplexed imaging results. Gene ontology (GO) analysis of DEGs enriched in tetrad CD8⁺ T cells revealed significant upregulation of pathways related to T-cell activation and T-cell-mediated cytotoxicity (Fig. 2I). Together, these findings indicate that tetrad formation associates with CD8⁺ T-cell effector activation while restraining terminal differentiation and dysfunction.

### Tetrad formation depends on the ICOS-ICOSL pathway

As noted above, the ICOS-ICOSL pathway is essential for the efficacy of anti-CTLA-4 therapy, which uniquely expands ICOS⁺ Th1-like CD4⁺ T cells—a feature not observed with anti-PD-1 treatment. However, how ICOS and ICOSL-expressing cells are spatially organized within tumors remains poorly understood. We therefore investigated whether ICOS-ICOSL signaling contributes to tetrad formation or function. Spatial analysis revealed that ICOS (protein) and *Icosl* (RNA) expressions were highest within tetrad regions, intermediate at 50 and 100 μm from tetrads, and lowest beyond 100 μm (Fig. 3A-C), indicating a tight spatial association between ICOS-ICOSL activity and tetrads.

**Figure 3:**
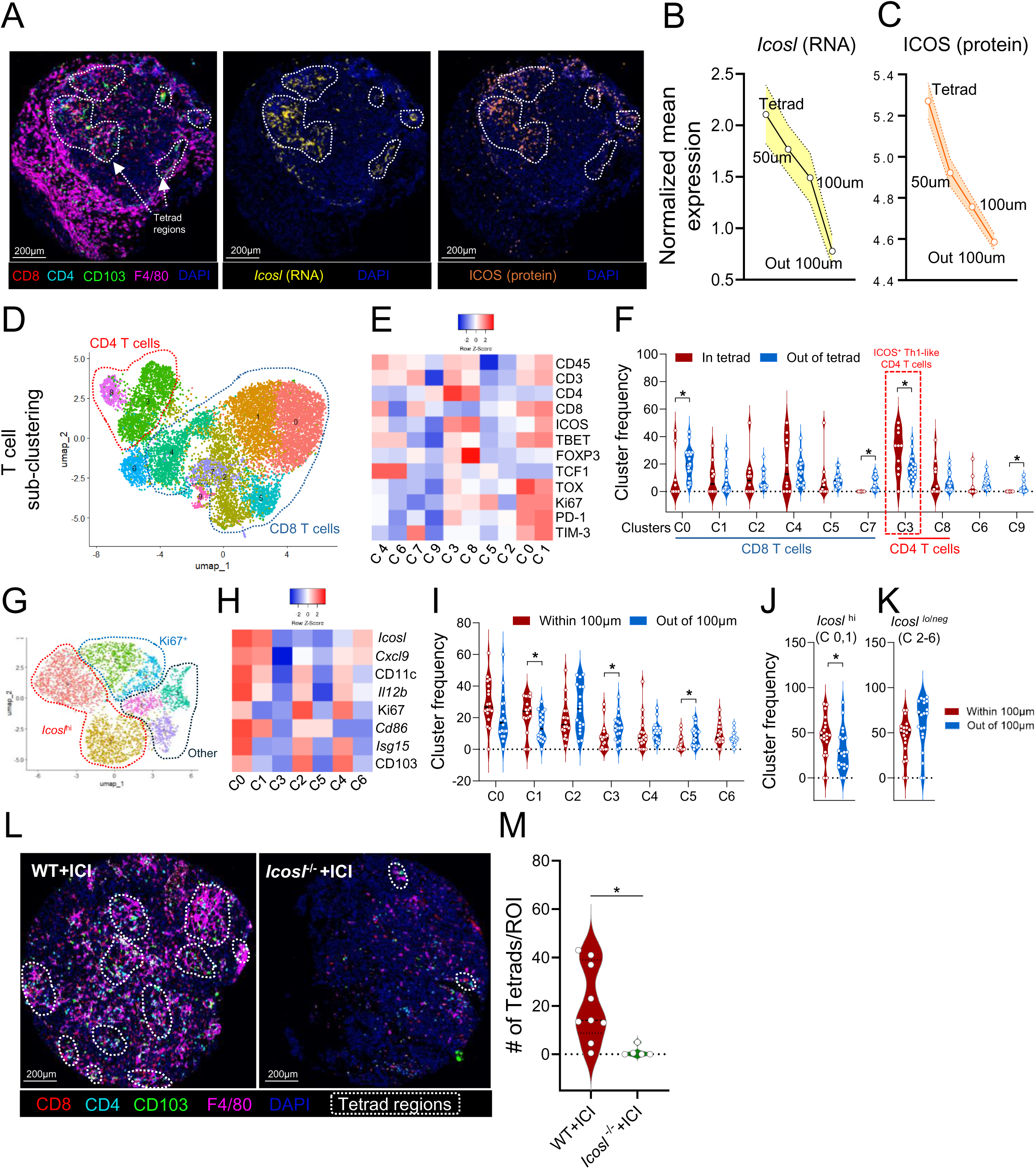
Tetrad formation depends on the ICOS-ICOSL pathway. **(A)** Representative COMET images highlighting tetrad regions, ICOS (orange) and *Icosl* (yellow) expressions. **(B-C)** Distribution of normalized mean expression of *Icosl* **(B)** and ICOS **(C)** in tetrads, within 50-μm and 100-μm neighborhoods, and outside of 100-μm. (n = 18-20 biological replicates). **(D-E)** UMAP **(D)** and heatmap **(E)** highlighting different T-cell sub-clusters and marker expression. **(F)** CD8^+^ and CD4^+^ T-cell sub-clusters frequency in tetrad versus outside of tetrads. (n = 10-20 biological replicates). (\**p*<0.05, unpaired T-test). Values are mean ± SEM. **(G-H)** UMAP **(G)** and heatmap **(H)** highlighting different cDC1 sub-clusters and marker expression. **(I-K)** cDC1 sub-clusters frequency **(I)**, *Icosl*^hi^ **(J)** and *Icosl*^lo/neg^ **(K)** clusters within 100-μm of tetrads versus outside of 100-μm of tetrads. (n = 18 biological replicates). (\**p*<0.05, unpaired T-test). Values are mean ± SEM. **(L)** Representative COMET images of day 12 B16F10 tumors isolated from wildtype or *Icosl*^−/−^ mice treated with two doses of anti-CTLA-4 and anti-PD-1. Tetrad regions are highlighted in white. **(M)** tetrad quantification from (L). (n = 5-9 biological replicates). (\**p*<0.05, unpaired T-test). Values are mean ± SEM.

To further define T-cell subsets engaged in these interactions, we subclustered T cells based on expression of TCF1, TOX, PD-1, TIM-3, T-BET, ICOS and other markers. We identified two CD4⁺ and six CD8⁺ T-cell clusters (Fig. 3D,E). Notably, cluster C3—characterized as ICOS⁺ T-BET⁺ Th1-like CD4⁺ T cells—was significantly enriched within tetrads compared with regions outside tetrads (Fig. 3F). In contrast, two CD8⁺ T-cell clusters were preferentially localized outside tetrads: cluster C0, expressing high levels of inhibitory receptors TIM-3, PD-1, and TOX, and cluster C7, defined by elevated TIM-3 (Fig. 3F), consistent with our earlier observations. The ICOS⁺ Th1-like CD4⁺ T-cell cluster (C3) exhibited a clear distance-dependent diffusion pattern—maximal within tetrads, intermediate at 50 and 100 μm, and lowest beyond 100 μm (Fig. S3A). Conversely, the terminally differentiated-like CD8⁺ cluster C0 showed the opposite gradient distribution, being most depleted within tetrads and progressively enriched with increasing distance (Fig. S3B). Together, these findings suggest that ICOS⁺ Th1-like CD4⁺ T cells are spatially organized around tetrads, whereas dysfunctional CD8⁺ T cells accumulate outside these regions.

We next subclustered cDC1s and identified seven different clusters, two of which (C0 and C1) exhibited high *Icosl* expression (Fig. 3G,H). These *Icosl*^high^ clusters were significantly enriched within tetrad regions (within 100 μm) compared to regions outside tetrads (Fig. 3I-K), although they did not display a pronounced distance-diffusion gradient (Fig. S3C). Macrophages showed a similar enrichment trend, but differences across clusters were not statistically significant (Fig. S3D-F). Single-cell RNA-seq analysis further revealed that *Icos* expression was highest in regulatory and non-regulatory CD4⁺ T cells and intermediate in CD8⁺ T cells (Fig. S3G), whereas *Icosl* expression was high in cDC1s (Fig. S3H). Together, these data indicate that ICOS⁺ Th1-like CD4⁺ T cells and ICOSL⁺ cDC1s are preferentially localized within tetrad regions, suggesting a key role for this pathway in the establishment and/or function of these multicellular structures.

To assess the functional relevance of ICOSL on cDC1s, we generated primary cDC1s from *Icosl*^+/+^ and *Icosl*^−/−^ bone marrow progenitors using a well-established differentiation protocol (24, 25) and adoptively transferred them into B16F10 tumor-bearing mice with or without anti-CTLA-4 treatment (Fig. S4A). Transfer of cDC1s alone, regardless of ICOSL expression, minimally affected tumor growth compared to controls. Strikingly, however, *Icosl*^+/+^ cDC1s combined with anti-CTLA-4 therapy eradicated 50% of tumors and significantly prolonged survival relative to both *Icosl*^−/−^ cDC1 + anti-CTLA-4 and anti-CTLA-4 monotherapy groups. These results demonstrate that ICOSL expression on cDC1s is functionally required for maximal antitumor efficacy of anti-CTLA-4 therapy.

Finally, to test whether the ICOS-ICOSL pathway contributes to tetrad assembly, we compared tetrad abundance in *Icosl*^−/−^ and wild-type mice bearing B16F10 tumors treated with ICI as in Fig. 1. Wild-type tumors exhibited significantly higher tetrad density compared to *Icosl*^−/−^ counterparts (Fig. 3L,M), indicating that ICOS-ICOSL signaling is required for tetrad formation either directly or indirectly. Collectively, these findings establish a mechanistic link between ICOS-ICOSL signaling and tetrad organization, highlighting this pathway as a critical determinant of early immune priming and therapeutic response.

### Tetrad-associated macrophages display an interferon-γ-responsive signature linked to reduced CD8⁺ T-cell dysfunction

Previous studies have shown that immunostimulatory macrophages correlate with favorable clinical outcomes in cancer patients ^30,31^. These macrophages are recruited by immunotherapy-activated T cells and are essential for therapeutic efficacy ^22^. Furthermore, both anti-CTLA-4 therapy and ICOS costimulation have been shown to reprogram macrophages toward an antitumor phenotype ^17,32^. However, the mechanisms by which these macrophages promote antitumor immunity and their spatial organization within tumors remain poorly understood. To test whether macrophages contribute to the assembly of cDC1-CD4-CD8 T-cell interactions, we depleted macrophages using a combination of anti-CSF-1R and anti-F4/80 antibodies. This approach efficiently decreased macrophage frequencies within tumors while preserving cDC1, CD4⁺, and CD8⁺ T-cell frequencies at day 12 post-tumor implantation (Fig. S5A,B). Macrophage depletion was then performed in conjunction with ICI treatment, as outlined in (Fig. 4A), and total macrophage numbers were markedly reduced per region of interest (ROI) (Fig. 4B). We next quantified cDC1-CD4-CD8 T-cell interactions independently of macrophage markers to assess triad formation in macrophage-depleted tumors. Strikingly, macrophage depletion significantly impaired triad formation (Fig. 4C), suggesting that macrophages orchestrate the assembly of APC-T-cell clusters such as tetrads. We note, however, that the depletion strategy employed here targets both immunostimulatory and immunosuppressive macrophages simultaneously, limiting the ability to ascribe this effect to a specific subset. The lack of tools for selective depletion of macrophage subpopulations remains a key challenge for the field.

**Figure 4:**
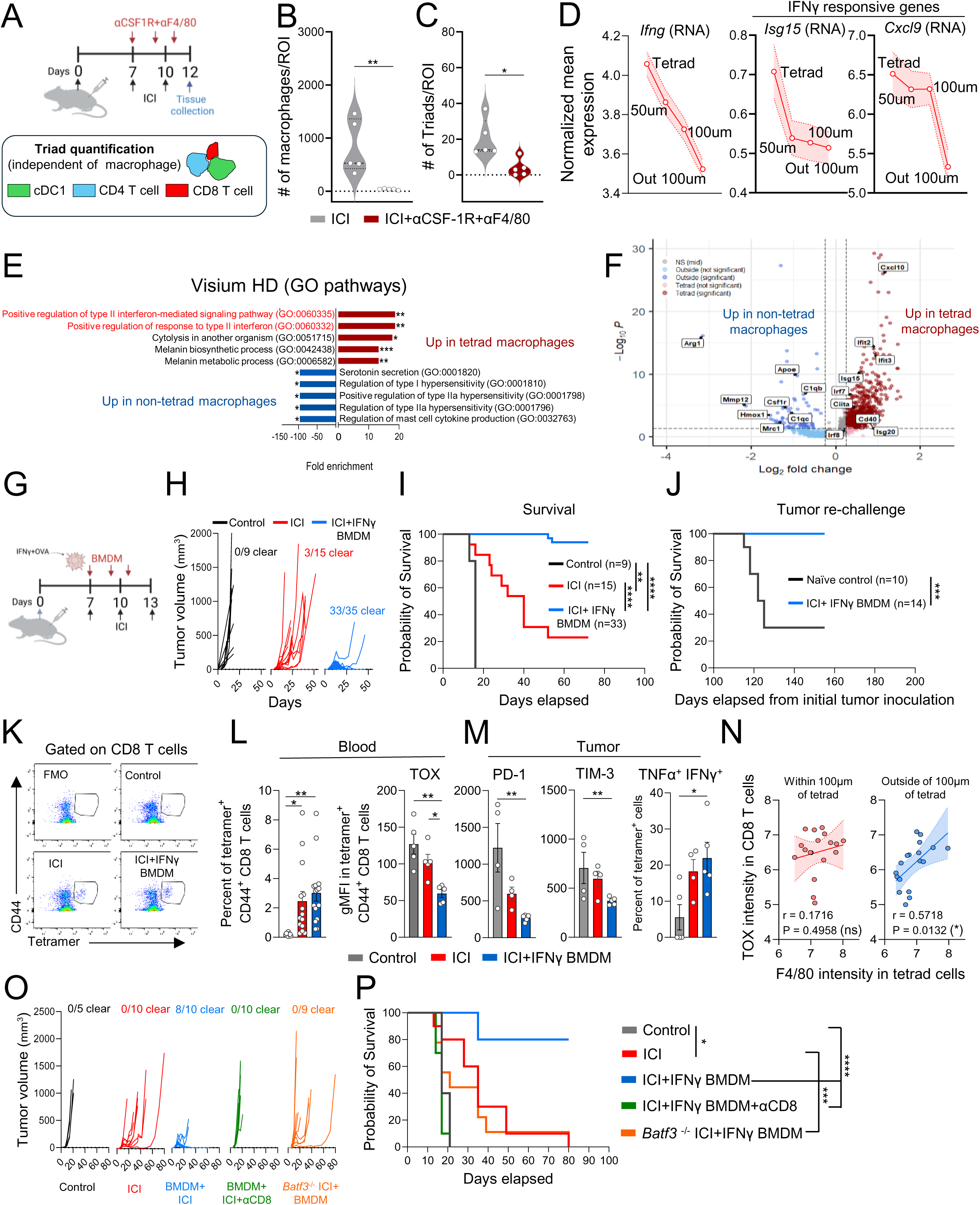
Tetrad-associated macrophages display an interferon-γ-responsive signature linked to reduced CD8⁺ T-cell dysfunction. **(A)** Schematic illustration of macrophage depletion experimental design and cDC1-CD4-CD8 triad quantification independent of macrophages. **(B-C)** Macrophages depletion efficiency **(B)** and cDC1-CD4-CD8 triad quantification between combination of anti-CTLA-4 and anti-PD-1 and anti-CTLA-4 and anti-PD-1 + anti-F4/80 +anti-CSF-1R **(C)**. (n = 5 biological replicates). (\**p*<0.05, \*\**p*<0.01, unpaired T-test). Values are mean ± SEM. **(D)** Distribution of normalized mean expression of *Ifng, Isg15 and Cxcl9* in tetrads, within 50-μm and 100-μm neighborhoods, and outside of 100-μm. (n = 18-20 biological replicates). **(E)** Top 5 GO terms enriched for Visium HD genes upregulated in tetrad (red) and non-tetrad (blue) macrophages. **(F)** Volcano plot for DEGs between tetrad (red) and non-tetrad (blue) macrophages. Statistically significantly DEGs (*p* < 0.05) are shown in red and blue, with select genes highlighted for reference. **(G-J)** *In vivo* adoptive transfer of IFNγ-stimulated bone marrow-derived macrophages (BMDMs) in combination with anti-CTLA-4 and anti-PD-1 as outlined in **(G)**. Outgrowth of B16F10-OVA tumors in B6 WT mice **(H)**. Data is pooled from 3 independent experiments (n = 9-35 mice per group). Kaplan-Meier survival curves of tumor-bearing mice are shown for mice challenged with primary tumors **(I)** (n = 9-35 mice per group), and re-challenged mice **(J)** (n = 10-14 mice per group). Data are shown as means ± SEM. Log-rank (Mantel-Cox) test was used to determine statistical significance for survival of mice (\*\**p*<0.01, \*\*\**p*<0.001, \*\*\*\**p*<0.0001). **(K-M)** Flow cytometry analysis of OVA₂₅₇-₂₆₄-specific tetramer^+^ CD44^+^ CD8^+^ T-cells showing tetramer^+^ cell frequencies and TOX expression in the blood (d13) **(L)** (n = 10-15 mice pooled from two independent experiments), and PD-1, TIM-3 and TNFα and IFNγ in tumor tetramer^+^ cells upon restimulation (d14) **(M)** (n= 5 mice per group). Significance is calculated by One-way ANOVA with Tukey’s multiple comparisons test. (\**p*<0.05, \*\**p*<0.01). Values are mean ± SEM. **(N)** linear regression best-fit lines displayed for TOX intensity in CD8^+^ T-cells with F4/80 intensity in tetrad member cells within 100-μm of tetrads versus outside of 100-μm of tetrads. (n = 18 biological replicates). **(O-P)** Outgrowth of B16F10-OVA tumors **(O)** and survival analysis **(P)** of *in vivo* adoptive transfer of IFNγ-stimulated BMDMs as in (G) into WT or *Batf3*^−/−^ mice or in combination with anti-CD8 depletion. (n = 5-10 mice per group). Data are shown as means ± SEM. Log-rank (Mantel-Cox) test was used to determine statistical significance for survival of mice (\**p*<0.05, \*\*\**p*<0.001, \*\*\*\**p*<0.0001).

We next sought to define the molecular and functional phenotype of tetrad-associated macrophages. Spatial RNAscope analysis revealed enrichment of *Ifng* and its downstream targets *Isg15* and *Cxcl9* within tetrad regions (Fig. 4D). Consistently, gene ontology analysis of differentially expressed genes (DEGs) from Visium HD data showed that the top two pathways enriched in tetrad macrophages were IFNγ-responsive signaling pathways (Fig. 4E). Among DEGs upregulated in tetrad-associated macrophages were interferon-inducible genes, including *Cxcl10*, *Irf7*, *Cd40*, *Isg20*, *Isg15*, and *Ciita*, whereas macrophages outside tetrads preferentially expressed immunosuppressive genes such as *Arg1*, *Mrc1*, *Apoe*, *C1qb*, and *Hmox1* (Fig. 4F). To test the functional relevance of this IFNγ signature, we adoptively transferred OVA-loaded IFNγ-stimulated bone marrow-derived macrophages (IFNγ-BMDMs) into B16F10-Ova tumor-bearing mice in combination with ICI (Fig. 4G). Strikingly, IFNγ-BMDM + ICI treatment eradicated 33 of 35 tumors, whereas ICI alone eliminated only 3 of 15 (Fig. 4H). This combination markedly prolonged survival compared with ICI alone or control groups (Fig. 4I). Moreover, mice that achieved complete tumor clearance resisted rechallenge, indicating durable T-cell memory formation (Fig. 4J). Phenotypic analysis of antigen-specific CD8⁺ T cells using tetramer staining (Fig. 4K) showed comparable frequencies between IFNγ-BMDM + ICI and ICI-alone groups in the blood; however, IFNγ-BMDM + ICI markedly reduced TOX expression in tetramer⁺ CD8⁺ T cells in the blood (Fig. 4L). Expression of the inhibitory receptors PD-1 and TIM-3 was also decreased in this group, whereas IFNγ and TNFα production was maintained in tumors (Fig. 4M), confirming preserved effector function. All analyses were performed prior to tumor rejection (day 13) to ensure that reduced inhibitory receptor expression was not confounded by tumor absence.

Spatially, macrophage abundance correlated strongly with TOX intensity outside tetrad regions but not within them (Fig. 4N), suggesting that IFNγ-responsive macrophages within tetrads may prevent CD8⁺ T-cell dysfunction. Finally, the therapeutic efficacy of IFNγ-BMDM + ICI required both CD8⁺ T cells and cDC1s: survival benefit was abrogated by CD8⁺ T-cell depletion or in *Batf3*⁻^/^⁻ mice lacking cDC1s (Fig. 4O,P). Collectively, these findings demonstrate that IFNγ-responsive macrophages are spatially enriched within tetrads, prevent CD8⁺ T-cell dysfunction, and are essential for durable antitumor immunity.

### Intratumoral tetrads predict response to dual checkpoint blockade in bladder cancer patients

We next asked whether tetrads could predict response to immunotherapy in humans. We analyzed tumor specimens from patients with urothelial carcinoma treated with dual checkpoint blockade—Durvalumab (anti-PD-L1) and Tremelimumab (anti-CTLA-4)—in a neoadjuvant clinical trial (27). Tumor samples were collected pre- and post-treatment (Fig. 5A). The cohort included nine patients, five responders and four non-responders based on pathologic assessment. Multiplexed imaging using CODEX was performed with a 65-marker panel, enabling phenotyping of over 5.7 million cells. Following major immune cell populations identification (Fig. S6A), we subclustered APCs and T cells to define cDC1s (BATF3⁺, HLA-DR^hi^, CD11c^int/lo^, BCL6⁺) (Fig. S6B) and CD4⁺ and CD8⁺ T-cell subsets (Fig. S6C). Using custom spatial algorithms, we quantified tetrads (macrophage-cDC1-CD4-CD8 interactions), triads (cDC1-CD4-CD8 without macrophages), and dyads (CD8-cDC1 or CD4-cDC1 pairs excluding macrophages). Strikingly, tetrad frequency, but not triads or dyads, was significantly higher in responders than in non-responders after treatment (Fig. 5B-E), indicating that tetrads are associated with pathologic response to dual immune checkpoint inhibition (ICI). Responders also exhibited higher baseline tetrad frequency than non-responders, suggesting that while ICI (likely via anti-CTLA-4) can induce tetrad formation, pre-existing tetrads may predict therapeutic responsiveness—consistent with our murine findings (Figs. 1, S1). Unexpectedly, tertiary lymphoid structure (TLS) density did not correlate with pathologic response in this cohort (Fig. 5F). However, all 9 patients analyzed were TLS high, and TLS abundance predicted response only in the broader clinical trial cohort (27). Moreover, tetrad density did not correlate with TLS density (Fig. 5G), implying that tetrads represent distinct microanatomical units, though we cannot exclude the possibility that tetrads may precede TLS formation.

**Figure 5:**
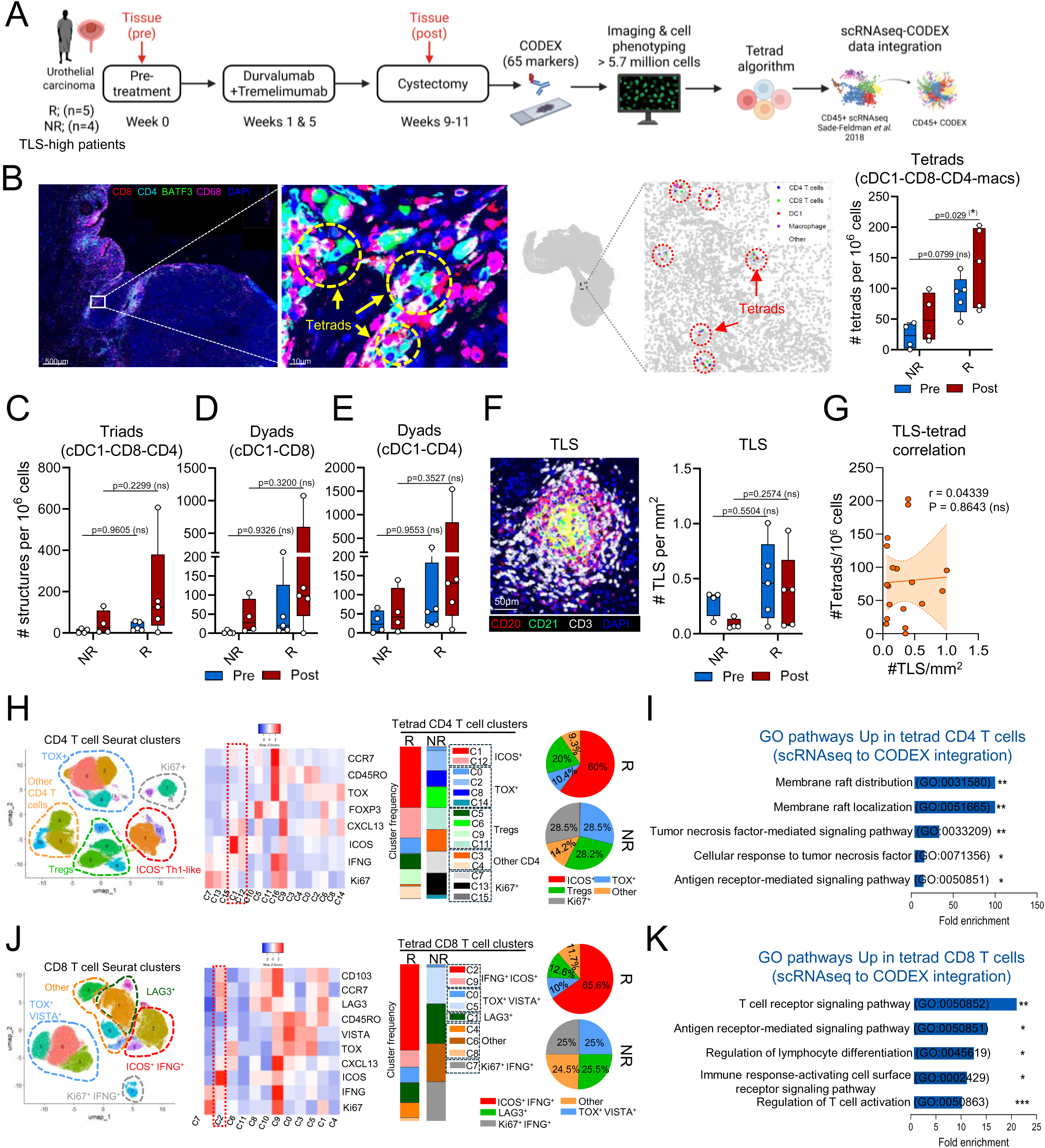
Intratumoral tetrads predict response to dual checkpoint blockade in bladder cancer patients. **(A)** Schematics illustrating the sequential immunofluorescence (CODEX) pipeline analysis for major cell phenotyping, tetrad identification and scRNAseq to CODEX data integration in human bladder cancer patients. **(B)** Tetrad quantification between responders and non-responders pre and post treatment. Significance is calculated by Two-way ANOVA of grouped R and NR with Šídák’s multiple comparisons test. (\**p*<0.05). Values are mean ± SEM. **(C-E)** Quantification of triads **(C)**, cDC1-CD8 dyads **(D)** and cDC1-CD4 dyads **(E)** in responders and non-responders pre and post treatment. Significance is calculated by Two-way ANOVA of grouped R and NR with Šídák’s multiple comparisons test. Values are mean ± SEM. **(F-G)** TLS abundance **(F)** and TLS/tetrad correlation **(G)** in responders and non-responders pre and post treatment. Significance is calculated by Two-way ANOVA of grouped R and NR with Šídák’s multiple comparisons test. Values are mean ± SEM. Linear regression best-fit lines displayed for TLS/tetrad correlation. **(H-K)** Frequency distribution of CD4⁺ T-cell subclusters **(H)** and CD8⁺ T-cell subclusters **(J)** in tetrad responders versus non-responders. Gene ontology pathway analyses for CD4⁺ T cells **(I)** and CD8⁺ T cells **(K)** were generated using differentially expressed genes derived from RNA integration.

Phenotypic analysis revealed that in responders, most tetrad-associated CD4⁺ T cells were ICOS⁺ Th1-like (60%), whereas these cells were nearly absent in non-responder tetrads (Fig. 5H), mirroring our mouse data. Similarly, tetrad-associated CD8⁺ T cells in responders were predominantly ICOS⁺ IFNγ⁺ (>65%) (Fig. 5J). To further interrogate functional programs, we integrated single-cell RNA-seq data from CD45⁺ tumor-infiltrating cells (28) into our CODEX dataset using a custom multimodal integration algorithm to infer gene expression spatially. Gene ontology analysis revealed that both tetrad-associated CD4⁺ (Figs. 5I, S6D) and CD8⁺ (Figs. 5K, S6E) T cells upregulated pathways involved in T-cell priming and activation compared to their non-tetrad counterparts, consistent with our murine observations. Tetrad-associated macrophages also upregulated IFNγ-responsive genes including CXCL9, CXCL10, and IFIT1 (Figs. S6F,G), paralleling the IFNγ-driven macrophage programs observed in mice. Together, these findings identify tetrads as spatial immune structures that mirror murine counterparts and predict clinical response to dual checkpoint blockade in bladder cancer.

## Discussion

In this study, we identify a multicellular assembly composed of cDC1, CD4⁺ T cells, CD8⁺ T cells, and macrophages—termed ‘tetrads’—that appears to coordinate antitumor effector function during immune-checkpoint blockade and may facilitate more effective priming of cytotoxic T cells. While intratumoral immune cell aggregates such as dyads ^33^, triads ^7,8^, and tertiary lymphoid structures ^34,35^ have been associated with response to ICIs, the mechanisms that guide the assembly of these structures remain incompletely understood. By characterizing tetrads and implicating ICOS-ICOSL signaling in their formation, our findings provide a framework for how spatially coordinated immune interactions may contribute to therapeutic response.

The spatial organization of tetrads appears to be closely linked to the unique biological effects of CTLA-4 and PD-1 blockade. CTLA-4 is rapidly upregulated following TCR engagement and attenuates early T-cell activation by outcompeting CD28 for the B7 ligands CD80 and CD86 ^36^. PD-1 on the other hand is induced later during T-cell activation in peripheral tissues and limits T-cell activity upon engagement with PD-L1 and PD-L2 ^37,38^. Prior studies, including our own, have shown that anti-CTLA-4 and anti-PD-1 therapies rescue T-cell activation through temporally and mechanistically distinct mechanisms ^1,10^. In particular, anti-CTLA-4 rescues T cell activation early during the priming phase ^10^, and selectively expands ICOS⁺ Th1-like CD4 effector T cells in mice and humans, a phenomenon not observed with anti-PD-1 therapy ^9^. The ICOS-ICOSL pathway has been shown to play a crucial role in the antitumor effects of anti-CTLA-4 and requires macrophages ^17^, yet how this axis influences multicellular immune organization has remained unclear. In this study, we report that tetrad formation is dependent on ICOS-ICOSL interactions and is characterized by the preferential enrichment of ICOS⁺ Th1-like CD4 T cells and ICOSL^high^ cDC1s within these structures. The elevated expression of ICOS on CD4^+^ T-cells may confer a selective advantage for engaging ICOSL on cDC1s, thereby initiating and/or sustaining tetrad assembly and enabling efficient CD8⁺ T-cell activation, although we cannot exclude possible involvement of other cell types in this pathway. Such early organizational cues may occur independently of more complex tertiary lymphoid structures that involve CD4⁺ T cells, CD8⁺ T cells, B cells, DCs and stromal cells, though we cannot exclude tetrads as potential precursors of TLS formation. These spatial distinctions help explain how anti-CTLA-4 and anti-PD-1 therapies act in complementary rather than redundant roles to shape antitumor immunity: anti-CTLA-4 enhances early T-cell activation during priming, whereas anti-PD-1 rescues T-cell activation post-priming, consistent with the superior clinical benefit of combination therapy in melanoma patients compared to monotherapies ^18,19^.

Although anti-CTLA-4 consistently promoted tetrad formation in both mouse and human tumors, baseline tetrad levels were higher in responder bladder cancer patients (Fig. 5B) and in murine melanoma tumors that display complete response to ICI such as mLama (Fig. S1). This suggests that tetrad formation is not exclusively induced by anti-CTLA-4. One possibility is that strong tumor antigens that can elicit potent immune responses may drive spontaneous tetrad assembly in highly immunogenic tumors, whereas anti-CTLA-4 may lower the activation threshold by reducing the requirement for strong MHC-peptide/TCR interactions, thereby enabling tetrad formation against weaker antigens. Together, these observations support a model in which tetrad formation reflects a convergence of antigenic strength and ICOS-dependent T-cell licensing, offering a mechanistic framework for understanding the complementary actions of CTLA-4 and PD-1 blockade, while introducing a novel APC-T cell organizational unit that may represent a more generalizable principle of immune coordination beyond checkpoint inhibition.

A defining feature that distinguishes tetrads from previously described immune triads is the presence of macrophages as a fourth cell type. Although tumor-associated macrophages (TAMs) are often linked to poor prognosis and therapy resistance ^20^. Emerging evidence indicates that macrophages can also support antitumor immunity ^21,22^, yet the mechanisms by which immunostimulatory macrophages are organized within tumors remain unclear. Consistent with prior work showing the requirement for macrophages in the efficacy of CTLA-4 blockade and ICOS costimulation ^17,32^, we find that tetrad-associated macrophages exhibit an interferon-γ-responsive transcriptional signature characteristic of immunostimulatory states. Functionally, interferon-γ-stimulated macrophages, when combined with immune-checkpoint inhibition, were sufficient to reject melanoma tumors in a manner dependent on CD8 T cells and cDC1s, underscoring a cooperative macrophage-cDC1 axis that extends beyond canonical dyadic interactions. Notably, these macrophages also limited antigen-specific CD8^+^ T-cell terminal differentiation compared to checkpoint blockade alone, suggesting a role in sustaining productive priming and preventing T-cell dysfunction.

Tetrads also differ from triads in the specific immune subsets they incorporate. Recent reports indicate that triads involve mregDCs, T follicular helper-like CD4 T cells, and stem-like CD8^+^ T cells ^7,8^, whereas tetrads are enriched for ICOS⁺ Th1-like CD4 T cells, cDC1s, and effector-like CD8^+^ T cells in both mice and humans. These distinctions may reflect temporal differences in the intratumoral immune response, with tetrads representing an earlier priming-associated structure preferentially promoted by anti-CTLA-4. We therefore propose that tetrads and triads are largely distinct but not mutually exclusive and may act in complementary roles whereby tetrads generate effector-like CD8^+^ T-cell pools early during priming, and triads support stem-like CD8^+^ T-cell pools, with both structures contributing to the prevention of CD8^+^ T-cell terminal differentiation.

Together, these findings establish tetrads as a novel intratumoral immune microstructure that integrates cDC1s, CD4^+^ T-cell, cytotoxic T cells, and macrophages to coordinate effective antitumor immunity. Our work provides a spatial and temporal framework for understanding how checkpoint therapies act synergistically. The cooperative macrophage-cDC1 axis, together with the complementary roles of tetrads and triads, suggests that tumors deploy multiple multicellular structures to generate and sustain productive CD8⁺ T-cell responses. The presence of tetrads across species, their enrichment in responders, and their spontaneous emergence in immunogenic tumors indicate that tetrads reflect a general mode of immune coordination rather than a treatment-specific phenomenon. These insights may inform strategies to overcome resistance to immune-checkpoint inhibition, including approaches that amplify ICOS signaling or enhance macrophage-directed antitumor functions. Defining how such structures arise and how they can be therapeutically reinforced may enable interventions that harness multicellular organization as a determinant of effective antitumor immunity.

## Methods

### Patient samples

For details on the clinical trial, see Gao et al.^35^. Briefly, Patients with high-risk, cisplatin-ineligible urothelial carcinoma (UC) were enrolled onto a pilot, single-arm neoadjuvant clinical trial evaluating combined PD-L1 and CTLA-4 blockade (MD Anderson Cancer Center protocol 2016-0033; NCT02812420). The study was approved by the MDACC Institutional Review Board, and all participants provided written informed consent. The primary endpoint was safety and tolerability of durvalumab plus tremelimumab in the neoadjuvant setting, assessed at each clinical visit according to NCI CTCAE v4.03. Secondary endpoints included immunological and molecular responses to therapy and pathological responses at cystectomy. Patients in this cohort received durvalumab (1,500 mg) plus tremelimumab (75 mg) on weeks 1 and 5. Radical cystectomy with bilateral pelvic lymph-node dissection was performed between weeks 9 and 11. Peripheral blood and tumor samples were collected pre- and post-treatment under a correlative laboratory protocol (MDACC PA13-0291).

### Mice

C57BL/6J and B6.129S(C)-*Batf3*^tm1Kmm/J^ mice were purchased from The Jackson Laboratory. *Icosl*^−/−^ mice on the B6 background were obtained from Dr. Padmanee Sharma at The University of Texas MD Anderson Cancer Center (MDACC) and are also available at The Jackson Laboratory. All animal studies were conducted in an AAALAC-accredited MDACC barrier facility in accordance with protocols approved by the MDACC Institutional Animal Care and Use Committee (IACUC).

### Cell lines

The mouse melanoma cell line B16-F10 was obtained from Dr. Isaiah Fidler (MD Anderson Cancer Center, Houston, TX, USA). The cell lines underwent authentication through spectral karyotyping to detect other cell contamination, and regular mycoplasma testing was conducted. Murine B16-F10 melanoma cells expressing OVA (B16-OVA cells) were obtained from Dr. Stephanie Watowich at MDACC. The *Braf*^V600E^ *Cdkn2a*^−/−^ *Pten*^−/−^ YUMM1.7 engineered to express defined MHC-I (mLama4) and MHC-II (mItgb1) neoantigens (mLama) were obtained from Dr. Matthew Gubin at MDACC. B16-F10 cells were cultured in Dulbecco’s Modified Eagle’s Medium (DMEM) supplemented with 10% fetal bovine serum (FBS, Corning) and penicillin-streptomycin solution at 37C in a 5% CO2 humidified incubator. B16-OVA and mLama cells were cultured in RPMI-1640 medium supplemented with 10% fetal bovine serum (FBS, Corning) and penicillin-streptomycin and cultured with the same conditions as B16-F10 cells. Cell lines were routinely tested for mycobacterium contamination.

### *In vivo* murine tumor experiments

Mice were co-housed and acclimated to the animal facility for at least one week before experimentation. For all in vivo studies, 8-12-week-old female mice were injected subcutaneously in the right flank with 2.5 × 10^5^ B16-F10 or B16-OVA cells, or 7.5 × 10^5^ mLama cells per mouse. Animals were randomized before initiation of checkpoint blockade therapy. For multiplexed imaging experiments, mice received intraperitoneal anti-CTLA-4 (clone 9H10, BioXCell) and/or anti-PD-1 (clone RMP1-14, BioXCell) on days 7 and 9 after tumor implantation. Tumors were harvested on day 12 for fixation, processing, and imaging. Anti-CTLA-4 was administered at 200 μg per mouse for the initial dose and 100 μg for subsequent doses; anti-PD-1 was administered at 250 μg per mouse. Tumor growth was monitored using digital calipers, and tumor volume was calculated as: volume = (width)^2^ × (length) / 2, as previously described ^39^. At study endpoint, mice were euthanized by CO₂ inhalation followed by cervical dislocation.

### *In vivo* macrophage depletion

To deplete macrophages in vivo, B16-F10 tumor-bearing mice were treated intraperitoneally with anti-CSF1R (clone AFS98, BioXCell) and anti-F4/80 (clone CI:A3-1, BioXCell) on days 7, 9, and 11 after tumor implantation. Each depleting antibody was administered at 500 μg for the initial dose and 250 μg for subsequent doses. Depletion efficiency was assessed by flow cytometry and multiplexed imaging. For multiplexed imaging experiments, macrophage depletion was performed in combination with dual checkpoint blockade using anti-CTLA-4 and anti-PD-1, as described above.

### Primary macrophage generation and stimulation

Bone marrow-derived macrophages (BMDMs) were generated as previously described ^39^. Briefly, Bone marrow cells were isolated from the femurs and tibias of age-matched female C57BL/6J mice and cultured in complete RPMI medium (5 × 10^5^ cells ml^−1^) supplemented with 50 μM β-mercaptoethanol, 10 mM HEPES, and 1 mM MEM non-essential amino acids (all from Thermo Fisher Scientific). BMDMs were differentiated for 5-7 days in the presence of M-CSF (50 ng ml^−1^) and subsequently stimulated with recombinant mouse IFNγ (20 ng ml^−1^; BioLegend) and supplemented with whole-protein ovalbumin (100 μg ml^−1^; InvivoGen) for 24 h before adoptive transfer. Cells were detached using Accutase (STEMCELL Technologies), washed twice with PBS, and resuspended for injection.

### Macrophage adoptive transfer

After randomization, 1 × 10^6^ activated BMDMs were delivered intratumorally into B16-F10-OVA tumor-bearing C57BL/6J wild-type or *Batf3*^−/−^ mice on days 7, 9, and 11, in combination with dual CTLA-4 and PD-1 blockade (administered on days 7, 10, and 13). Tumor growth was monitored as described above. Survival endpoints were defined as death, moribund status, or tumor diameter ≥15 mm in any direction per IACUC guidelines. For tumor rechallenge experiments, tumor-cleared mice and naïve controls were injected with B16-F10-OVA cells 100 days post initial tumor challenge and as described above.

### *In vivo* CD8^+^ T-cell depletion

For CD8⁺ T-cell depletion, anti-CD8α (clone 2.43, Bio X Cell) was administered on day 6 (500 μg) and days 9 and 12 (250 μg) concomitantly with dual checkpoint blockade and BMDM transfer. Depletion efficiency was confirmed by measuring CD8⁺ T-cell frequency in peripheral blood.

### Primary cDC1 generation

Primary cDC1s were generated from bone marrow cells of C57BL/6J wild-type or *Icosl*^−/−^ mice as previously described, with minor modifications ^40,41^. Briefly, bone marrow cells were cultured in complete RPMI 1640 medium supplemented with FMS-like tyrosine kinase 3 ligand (FLT3L; 200 ng/ml) and granulocyte-macrophage colony-stimulating factor (GM-CSF; 5 ng/ml) at a density of 1.5 × 10□ cells/ml in a total volume of 10 ml. On day 5, cultures were supplemented with an additional 5 ml of complete medium. On day 9, non-adherent cells were collected and re-cultured in complete medium containing FLT3L (50 ng/ml) and GM-CSF (5 ng/ml) at a density of 3 × 10□ cells/ml. Non-adherent cells were harvested between days 16 and 18, and cDC1 purity (CD11c⁺ CD103⁺ CD24⁺ B220⁻ CD172α⁻) was assessed by flow cytometry, consistently exceeding 70%. ICOSL expression was confirmed to be absent in *Icosl*⁻^/^⁻ cDC1s and present in >95% of wild-type cDC1s as evaluated by flow cytometry.

### cDC1 *in vivo* adoptive transfer

For adoptive transfer experiments, 2 × 10□ primary cDC1s were stimulated overnight with poly(deoxyinosinic-deoxycytidylic) acid sodium salt (poly I:C; 20 μg/ml; Sigma-Aldrich) and subsequently injected intratumorally into B16F10 tumor-bearing mice on days 7 and 10, either alone or in combination with anti-CTLA-4 antibody treatment (administered on days 7, 10, 13, and 16). Tumor growth and survival were analyzed as described above.

### Tumor tissue processing, flow cytometry and tetramer staining

In macrophage adoptive transfer experiments, B16F10-OVA tumor-bearing mice were euthanized on day 14 post-tumor implantation, prior to tumor rejection in the BMDM + ICI group, to prevent loss of antigenic source during analysis of tumor-specific CD8⁺ T cells. Tumors were excised and enzymatically digested using Liberase TL (Roche) and DNase I (Roche) at 37 °C for 30 min with gentle agitation every 5 min. Digested tissues were passed through a 70-μm nylon cell strainer to generate single-cell suspensions. Murine whole blood samples were treated with red blood cell (RBC) lysis buffer (Sigma-Aldrich) to deplete RBCs and obtain single cell suspension. Flow cytometry was performed as previously described ^39^. Briefly, 3 × 10□ cells per mouse were stained with Live/Dead Fixable Ghost Dye (Tonbo Biosciences), followed by Fc receptor blocking (BioXCell) and surface antibody staining. Cells were subsequently fixed and permeabilized using the FoxP3 Fixation/Permeabilization Buffer Kit (Thermo Fisher Scientific) according to the manufacturer’s instructions and stained with intracellular antibodies. Fluorescence minus one (FMO) controls and single-color UltraComp eBeads (Invitrogen) were used as negative and positive controls, respectively. Data were acquired on an LSRFortessa X-20 flow cytometer (BD Biosciences).

For intracellular cytokine staining, tumor-derived single-cell suspensions were stimulated with Cell Activation Cocktail (BioLegend) for 4 h to allow accumulation of intracellular cytokines, following the manufacturer’s protocol. After surface marker staining, cells were fixed, permeabilized, and stained with antibodies against IFN-γ, TNF-α, and other intracellular markers listed in the Supplementary Materials. Antigen-specific CD8⁺ T cells were identified using an OVA₂₅₇-₂₆₄-specific tetramer (Baylor College of Medicine Tetramer Core Facility) and were gated as singlets, live CD45⁺ CD3⁺ CD8⁺ CD44⁺ tetramer⁺ cells. Flow cytometry data were analyzed using FlowJo software v10 (Tree Star). All antibodies are listed in supplementary tables.

### RNAScope with sequential immunofluorescence staining (COMET) of mouse TMAs

Formalin-fixed mouse tumors were processed for paraffin embedding (FFPE), sectioned at 5 μm, and stained with hematoxylin and eosin (H&E). Tissue microarrays (TMAs) were constructed by randomly selecting two 1-mm-diameter regions of interest (ROIs) per tumor based on H&E images. To minimize RNase contamination, all tissue processing steps were performed using nuclease-free reagents, and work surfaces were treated with RNaseZap (Invitrogen, AM9784) followed by 70% ethanol. FFPE sections were baked at 60 °C for 1 h, deparaffinized in xylene (Sigma, 247642; 5 min, three changes), and rehydrated through a graded ethanol series (100%, 95%, 70%, 50%, and 30%) before equilibration in PBS. Antigen retrieval was performed using an EZ Retriever System (Biogenex, v3) at 105 °C for 15 min in AR 2 Elegance buffer (Biogenex, HK547-XAK; pH 9). Slides were rinsed in PBS and subsequently loaded onto the COMET platform (Lunaphore, NSP/1.0).

RNAscope HiPlex Pro assays (Advanced Cell Diagnostics, 322070) were performed according to the manufacturer’s protocol, with probes diluted 1:50 in probe dilution buffer. Sections underwent protease-free pretreatment, followed by probe hybridization, signal amplification, and three iterative rounds of fluorescent probe detection and cleavage. Following RNAscope processing, sequential immunofluorescence (seqIF) was performed using unconjugated primary antibodies and corresponding secondary antibodies. Antibody order, incubation times, and imaging parameters were optimized empirically. After antibody elution, a second seqIF cycle was conducted, enabling detection of up to 12 RNA targets and 20 protein markers per section. All antibodies and RNA probes are listed in supplementary tables.

### COMET data preprocessing and cell phenotyping

Whole-slide COMET OME-TIFF tumor core images were preprocessed by subtracting the background using the Lunaphore Horizon software, then by splitting each image into non-overlapping 1024 × 1024 fields of view (FOVs), from which individual TIFF files for every channel were extracted. Cell segmentation was performed using the Mesmer algorithm ^42^, with DAPI used to identify nuclei and the sum of all non-nuclear markers used to define whole-cell boundaries. After segmentation, we also computed morphological features per cell, including area, perimeter, eccentricity, major and minor axis lengths, and centroid coordinates for downstream analysis. Image quality control included removal of FOVs that have less than 10 cells per FOV. Pixel-level phenotyping was then carried out using Pixie^24^, which clusters multi-channel pixel intensities; lineage markers including CD45, CD103, CD11c, CD4, CD3e, CD11b, CD8, B220 and F4/80 were used to interpret pixel clusters. For each segmented cell, we quantified the number of pixels clusters and normalized these counts by cell area to generate per-cell features. Cells were subsequently clustered using a self-organizing map (SOM)^24,43,44^, and metaclusters were identified using consensus clustering. Final major cell type annotations were assigned based on the lineage marker enrichment patterns of the metaclusters.

### CODEX staining of human tumor sections

CODEX staining assays were performed according to the manufacturer’s protocol. Formalin-fixed, paraffin-embedded (FFPE) human tumor tissues were sectioned at 4 μm, mounted onto glass slides, and stored at 4 °C until use. Purified, BSA-free primary antibodies were obtained from commercial vendors and are listed in the Supplementary Tables. Commercially available barcoded antibodies, reporter reagents, and antibody conjugation kits were purchased from Akoya Biosciences. Unconjugated antibodies were conjugated in-house according to the manufacturer’s recommendations, as previously described ^45^. CODEX staining protocols were validated using human tonsil tissue and control tumor sections. Antigen retrieval was performed using Tris-EDTA buffer (pH 9.0). Antibodies were incubated either overnight at 4 °C or for 3 h at room temperature, depending on the optimized staining conditions. Multiplexed imaging was performed using the PhenoCycler™-Fusion system (Akoya Biosciences).

### CODEX analysis of human tumor sections

Segmentation was performed using Visiopharm (20250.08.2) Cell Detection-AI (Fluorescence) algorithm. Cell clustering and phenotyping was performed using the Leiden graph clustering algorithm to identify major cell populations. Minor cell populations such as CD4^+^ and CD8^+^ T-cell populations were identified by sub-clustering CD3^+^ T-cells using phenotypical and functional markers as outlined in figures and results section. cDC1 cells were identified by sub-clustering the 75^th^ percentile (upper quartile) of HLA-DR expression in all annotated cells (high HLA-DR cells) where DC, macrophage and B cell-specific markers were used. cDC1 cluster was defined as: BATF3^+^, HLA-DR ^high^,CD1c ^low^, CD11c ^int/low^, BCL6^+^, IDO1^+^, CD68/14 ^neg^, CD20 ^neg^.

### Sub-clustering in COMET/RNAscope and CODEX Single-Cell Data

To resolve fine-grained minor cell states across our different multiplexed datasets, we performed sub-clustering of major cell types using the standard Seurat clustering workflow. Following cell segmentation and major cell type annotation as described above, the top 40 variable features were identified, and expression values were scaled prior to principal component analysis (PCA) using the top 30 principal components. A graph-based clustering was applied to identify distinct subclusters unique to each major cell population. Clusters were annotated based on co-expression of key phenotypic and functional markers and UMAP feature overlays and scaled expression heatmaps were used to confirm marker specificity. Heatmaps were generated using the publicly available tool http://www2.heatmapper.ca/expression/.

### Tetrad, triad and dyad identification algorithms

For mouse COMET datasets, we leveraged the subcellular spatial resolution of the COMET platform to infer cell-cell interactions based on direct physical contact. Interactions were defined by contiguous whole-cell polygon masks derived from inferred membrane boundaries, such that two cells were considered interacting when their mask borders were directly touching or connected. To identify immune tetrad structures, we developed a custom algorithm defining a tetrad as a cDC1 physically interacting with at least one CD4⁺ T cell and one CD8⁺ T cell, with all three cell types additionally expressing macrophage markers (F4/80 or CD64) above the first quartile of expression. This criterion was based on qualitative and quantitative observations from COMET images showing that activated macrophages exhibit a larger, more spread morphology compared to other immune cell types, resulting in consistent detection of macrophage marker signal across interacting neighboring cells. Thus, shared macrophage marker expression across the interacting cDC1, CD4⁺ T cell, and CD8⁺ T cell was used to infer the presence of macrophages participating in tetrad formation. Tetrads were quantified by counting the number of unique cDC1 cells participating in tetrad structures, thereby avoiding multiple counting of the same structure. Triads were defined as interacting cDC1-CD4⁺-CD8⁺ T-cell assemblies in the absence of macrophages, as indicated by macrophage marker expression (F4/80 or CD64) below the first quartile in all interacting cells. In macrophage depletion experiments, cDC1-CD4⁺-CD8⁺ interactions were quantified independent of macrophage presence by summing the number of triads and tetrads, thereby capturing total three-cell interaction events.

To identify tetrad structures in human CODEX samples, we developed a spatial proximity-based algorithm to detect four-cell assemblies composed of a cDC1, a CD4⁺ T cell, a CD8⁺ T cell, and a macrophage. Because our CODEX data lacked whole-cell polygon masks, tetrad identification was based on spatial distances rather than direct membrane contact. All DC1 cells were identified, and Euclidean distances were computed between each cDC1 cell and neighboring CD4⁺ T cells, CD8⁺ T cells, and macrophages. A tetrad was defined as a configuration in which a DC1 cell was located within a 20-µm radius of at least one CD4⁺ T cell, one CD8⁺ T cell, and one macrophage. For each cDC1 cell, all neighboring CD4⁺, CD8⁺, and macrophage cells within this search radius were enumerated. When all three partner cell types were simultaneously detected around a given cDC1 cell, the corresponding set of four cells was classified as a DC1-CD4⁺-CD8⁺-macrophage tetrad. To ensure that each cell contributed to at most one tetrad, a non-overlap constraint was applied, excluding any candidate tetrad containing a cell already assigned to a previously identified structure. This constraint ensured that tetrad counts reflected distinct cellular assemblies rather than repeated sampling of the same cells. For each TMA region, the total number of tetrads was calculated as the number of unique DC1 cells participating in valid four-cell tetrad arrangements. Triads were defined by the presence of all three cell types with the absence of macrophages, while Dyads were defined as two-cell assemblies composed of the specified pair of cell types, with exclusion of the remaining two cell types not included in the dyad.

### Identification of cells within 50 μm and 100 μm of cDC1-anchored tetrads

To characterize the cellular composition surrounding cDC1-CD4⁺-CD8⁺-macrophage tetrads, we quantified all cells located within defined spatial neighborhoods centered on tetrad-anchoring cDC1 cells. The centroid of each cDC1 participating in a tetrad was used to generate circular buffer regions with radii of 50 μm and 100 μm. These buffers defined spatial neighborhoods in which surrounding cells were queried. A cell was annotated as residing within a given neighborhood if its centroid fell within any cDC1-anchored buffer of the corresponding radius. Analyses were performed independently for the 50-μm and 100-μm neighborhoods to capture both immediate and extended cellular contexts surrounding tetrad structures.

### Tertiary Lymphoid Structure analysis in human CODEX data

Tertiary lymphoid structures (TLSs) were evaluated in FFPE tissue sections from patients with urothelial carcinoma (N = 9) at pre-treatment (n = 9) and post-treatment (n = 9) time points. Patients were classified as responders (n = 5) or non-responders (n = 4) based on pathological response criteria ^35^. Four-micron FFPE tissue sections were stained by immunohistochemistry using primary antibodies against CD20 (Dako, M075529) and CD3 (Dako, A0452), followed by appropriate secondary antibodies, peroxidase-conjugated avidin/biotin, and 3,3′-diaminobenzidine substrate (Leica Microsystems) for visualization. Slides were scanned using a ScanScope XT system (Leica Technologies) and analyzed by a pathologist using HALO v2.3 software (Indica Labs). Lymphoid aggregates and TLSs were manually identified and annotated. TLS density was calculated as the number of TLSs per mm² of annotated tissue area, as previously described ^34^.

### Spatial transcriptomic analysis of mouse tumors (Visium HD)

Visium HD spatial transcriptomics was performed as previously described ^28^. Briefly, FFPE B16F10 tumor samples were processed according to the Visium HD FFPE Tissue Preparation Handbook (CG000684) and Spatial Gene Expression Reagent Kits User Guide (CG000685) and sequenced on a NovaSeq 6000 SP-100 Xp instrument. Data preprocessing was performed using Space Ranger v3.0. A tissue microarray (TMA) containing 49 B16F10 tumor cores (biological replicates) was analyzed, with an adjacent TMA slide used for H&E staining and nucleus-based cell segmentation. To assign immune cell identities, we used a supervised reference-mapping approach in which a publicly available annotated CD45⁺ scRNA-seq dataset served as the reference ^29^. Shared transcriptional features (using 2,000 variable features) for each major immune cell population between the scRNA-seq reference and Visium HD CD45⁺ cells were identified and used to project Visium HD cells into the reference UMAP space. Each Visium HD cell was assigned the label of its closest matching scRNA-seq population, and these immune annotations were propagated to the full Visium HD dataset for downstream analyses.

Tetrad structures were identified by integrating inferred immune phenotypes with single-cell spatial coordinates. For each cDC1 cell, distances to neighboring CD4⁺ T cells, CD8⁺ T cells, and macrophages were computed, and a DC1 cell was classified as forming a tetrad when all three partner cell types were located within a 100-µm radius. Unique DC1-CD4⁺-CD8⁺-macrophage combinations were recorded using a non-overlap constraint to ensure that each cell contributed to at most one tetrad. Differential gene expression analyses comparing tetrad-associated versus non-associated CD4⁺ T cells, CD8⁺ T cells, and macrophages were performed using Seurat’s Wilcoxon rank-sum test. For each immune cell population, differentially expressed gene (DEG) lists were filtered to retain only genes specific to that population based on scRNA-seq data. Genes uniquely upregulated or downregulated in tetrad-associated cells compared with non-associated cells (p < 0.05) were used as input for biological process gene ontology analysis (http://geneontology.org/).

### Integration of CODEX with scRNA-seq Data

CODEX protein measurements were integrated with a CD45⁺ scRNA-seq reference dataset to infer RNA expressions for protein-resolved single cells. The CODEX dataset was first preprocessed where non-informative background and structural markers were removed, and each antibody channel was mapped to its corresponding gene symbol. Representative mappings included CD3e to *CD3E*, CD8 to *CD8A*, CD11c to *ITGAX*, PD-1 to *PDCD1*, TIM3 to *HAVCR2* and Ki67 to *MKI67*. Only protein channels with mapped genes present in the scRNA-seq reference were retained for downstream integration. Protein intensities were then normalized and scaled before integration.

The scRNA-seq reference ^46^ was processed by log-normalization, identification of variable genes, and principal component analysis. To align protein and RNA-based modalities, we used an anchor-based approach ^47,48^ which identifies pairs of cells across datasets that are predicted to represent the same underlying biological state. Shared protein-RNA features were projected into a joint low-dimensional space using reciprocal PCA, and anchors were defined as mutual nearest neighbors between CODEX and scRNA-seq cells in this shared space. After anchors were established, cell-type annotations from the scRNA-seq reference were transferred to CODEX cells using weighted predictions based on the local anchor neighborhoods. The same anchor weights were then used to impute RNA expression for genes not directly measured in the CODEX data. Feature imputation follows a generalized framework ^47^ in which predicted gene expression for a query cell is obtained by multiplying reference gene-expression values by anchor-derived weights, providing continuous estimates of transcript abundance. Differential gene expression and gene ontology analyses were performed using the same approach as for the Visium HD analysis.

### Quantification and statistical analysis

Statistical analyses were performed using Prism v10.6.1 (GraphPad Software) unless otherwise indicated. Details of statistical tests used for each experiment are provided in the corresponding figure legends. For comparisons between two groups in flow cytometry and spatial analyses, unpaired two-tailed Student’s t tests or other appropriate statistical tests were applied as specified in the figure legends. For mouse tetrad quantification, statistical significance was determined using one-way analysis of variance (ANOVA). Data are presented as mean ± SEM. A P value < 0.05 was considered statistically significant. Mice were randomized prior to group allocation.

## Acknowledgments

We thank all members of the Allison and Sharma labs for support throughout this project; Sequencing data was produced by the ATGC supported by the Core grant CA016672(ATGC) and NIH 1S10OD024977-01; We thank funding sources: Parker Institute for Cancer Immunotherapy and T.C. and Jeanette Hsu Foundation (PS, JPA), Ergon Foundation Award and Odyssey Fellowship (MC). We also thank Flow cytometry, histology and imaging cores at MD Anderson. We would like to give special thanks to the Immunotherapy platform for helping with human data and Visium HD.

## Author contributions

Conceptualization: MC, MA, PS and JPA.; methodology: MC, MA, SH, YX, AB, AC, MDM, SJ, SB, KHH, JW, JB; investigation: MC, MA, SH, MA, AB, AC, MDM; visualization: S.H. and N.G.; funding acquisition: JPA and PS; project administration: JPA and P.S.; supervision: MC, MA, JPA and P.S.; writing – original draft: MC, MA; writing – review and editing: MC, MA, SH, KHH, MG, XC, JJM, SW, JPA and PS.

## Competing interests

Authors declare no competing interests.

## Data and code availability

Sequencing and imaging processed data will be deposited in public repositories. Codes used in this study will be deposited on GitHub. Further details will be made available by the lead contacts on request.

## Author notes

Disclosures: P. Sharma reported “other” from Achelois, Adaptive Biotechnologies, Affini-T, Apricity, Asher Bio, BioAtla LLC, BioNTech, Candel Therapeutics, Catalio, Carisma, C-Reveal Therapeutics, Dragonfly Therapeutics, Earli, Inc., Enable Medicine, Glympse, Henlius/Hengenix, Hummingbird, ImaginAb, InterVenn Biosciences, JSL Health, LAVA Therapeutics, Lytix Biopharma, Marker Therapeutics, Oncolytics, PBM Capital, Phenomic AI, Polaris Pharma, Sporos, Time Bioventures, Trained Therapeutix Discovery, Two Bear Capital, Xilis, Inc., Akoya Biosciences, Osteologic Therapeutics, and Matrisome outside the submitted work. J.P. Allison reported personal fees from Achelois, Adaptive Biotechnologies, Akoya Biosciences, Apricity, Bectas, BioAtla, BioNTech, Candel Therapeutics, Dragonfly, Earli, Enable Medicine, Hummingbird, ImaginAb, Lava Therapeutics, Lytix, Marker, Osteologic, PBM Capital, Phenomic AI, Polaris Pharma, Time Bioventures, Trained Therapeutix, Two Bear Capital, and Venn Biosciences outside the submitted work. No other disclosures were reported.

## Extended Data Figures Legends

**Figure S1:**
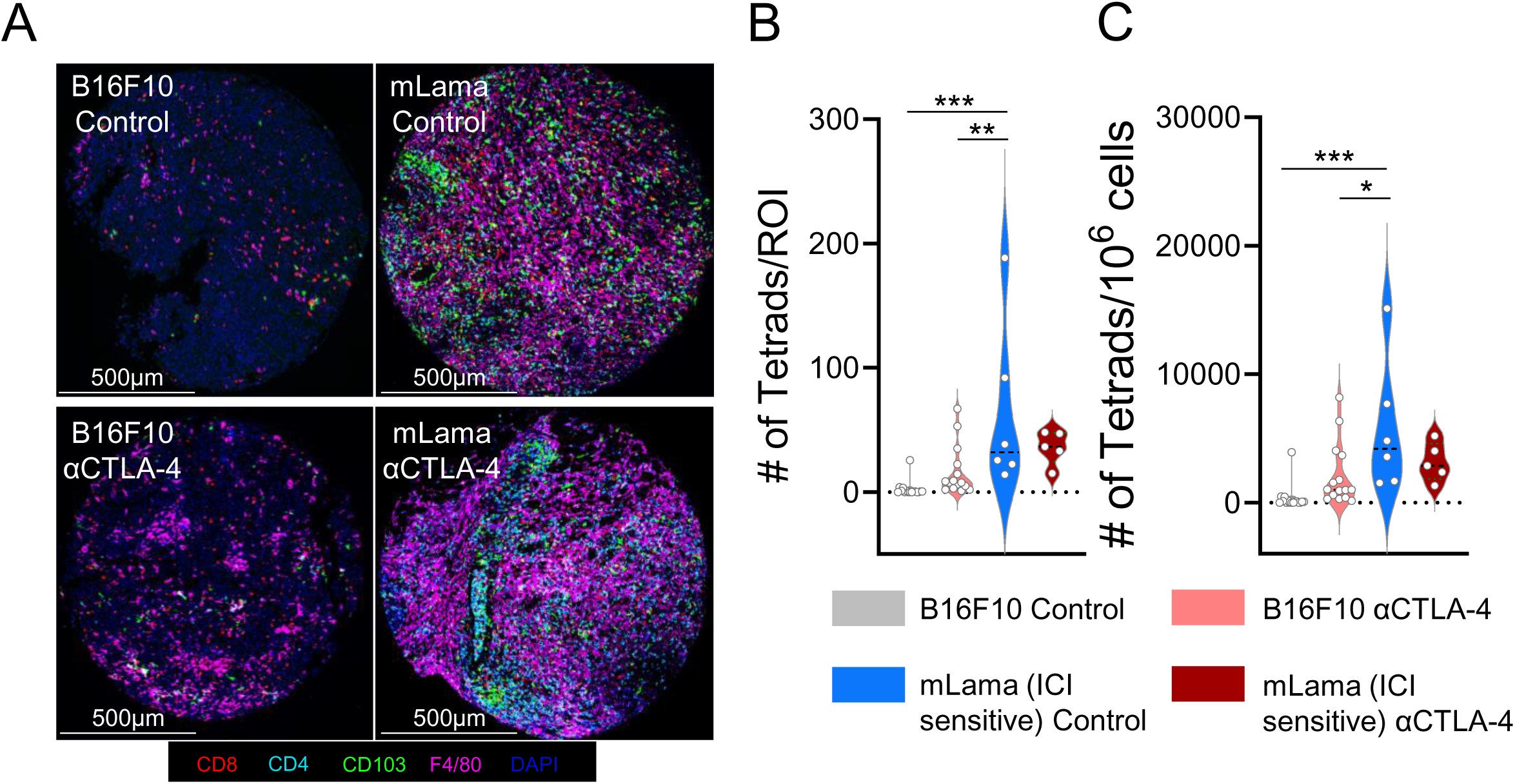
Comparison of tetrad frequencies in B16F10 and mLama tumors (related to figure 1) **(A)** Representative COMET images highlighting tetrad cells. **(B-C)** Tetrad quantification representing absolute numbers per ROI **(B)**, and numbers normalized to the total number of cells per each ROI **(C)**. (n = 5-16 biological replicates per group). Significance is calculated by One-way ANOVA with Tukey’s multiple comparisons test. (\**p*<0.05, \*\**p*<0.01, \*\*\**p*<0.001). Values are mean ± SEM.

**Figure S2:**
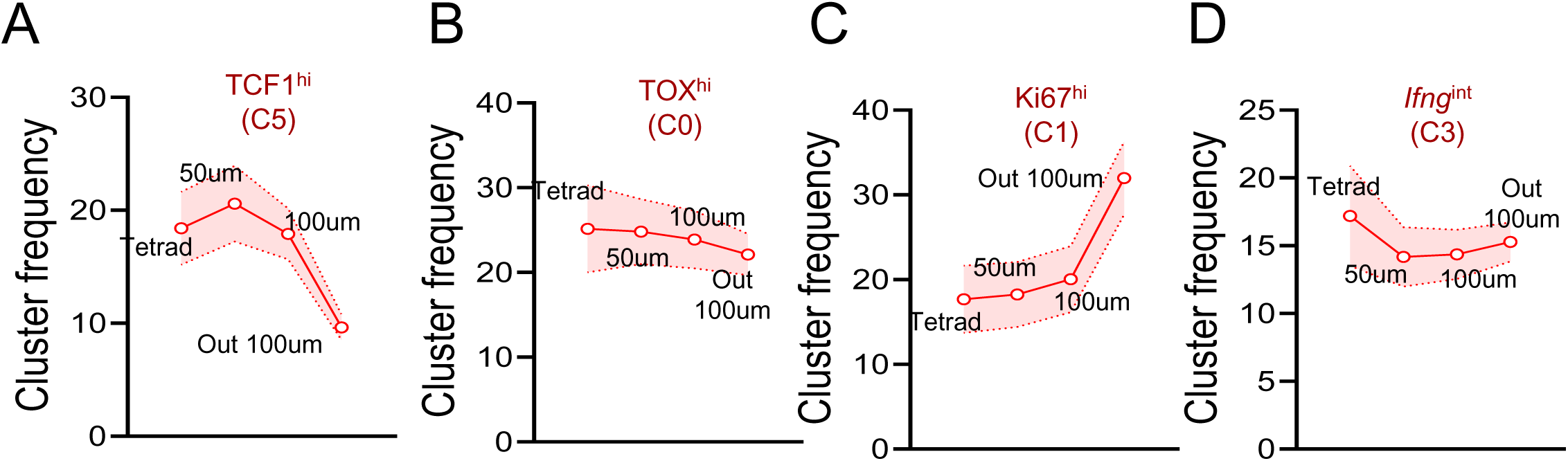
Spatial distribution of CD8+ T-cell sub-clusters in mouse B16F10 tumors (related to figure 2) **(A-D)** C5 (TCF1^hi^) **(A),** C0 (TOX^hi^) **(B),** C1 (Ki67^hi^) **(C)** and C3 (*Ifng*^hi^) **(D)** CD8^+^ T-cell sub-cluster frequency distribution in tetrads, within 50-μm and 100-μm neighborhoods, and outside of 100-μm. (n = 18 biological replicates)

**Figure S3:**
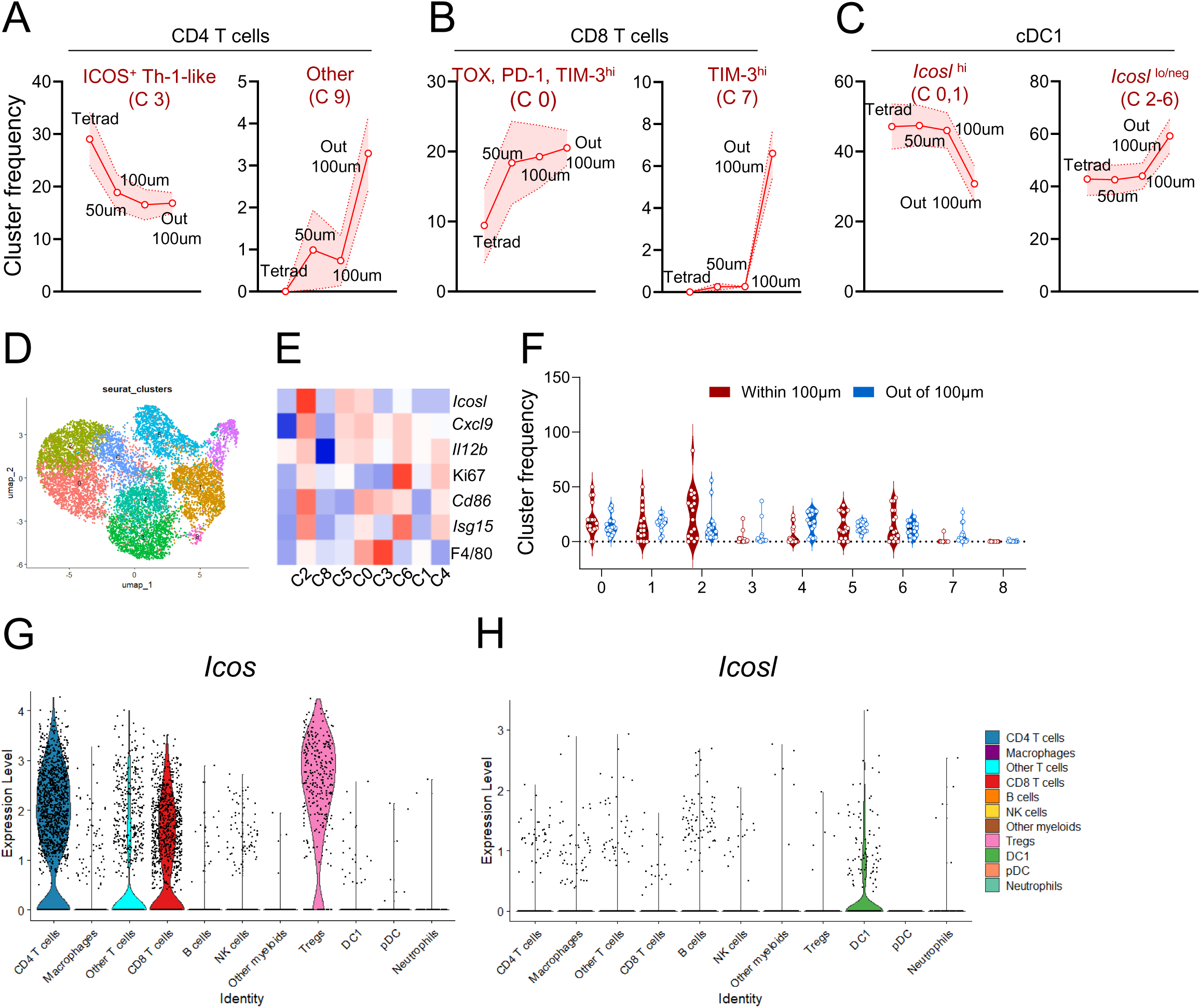
Tetrad formation depends on the ICOS-ICOSL pathway (related to figure 3) **(A-C)** Cluster frequency distribution of CD4^+^ T-cells **(A)**, CD8^+^ T-cells **(B)** and cDC1 **(C)** in tetrads, within 50-μm and 100-μm neighborhoods, and outside of 100-μm. **(D-F)** Macrophage sub-clusters shown in UMAP **(D)**, heatmap **(E)** and cluster frequency **(D)** within 100-μm of tetrads versus outside of 100-μm of tetrads. (n = 18 biological replicates). (unpaired T-test). Values are mean ± SEM. **(G-H)** Transcript expression of *Icos* **(G)** and *Icosl* **(H)** in intratumoral immune cell populations from scRNAseq data.

**Figure S4:**
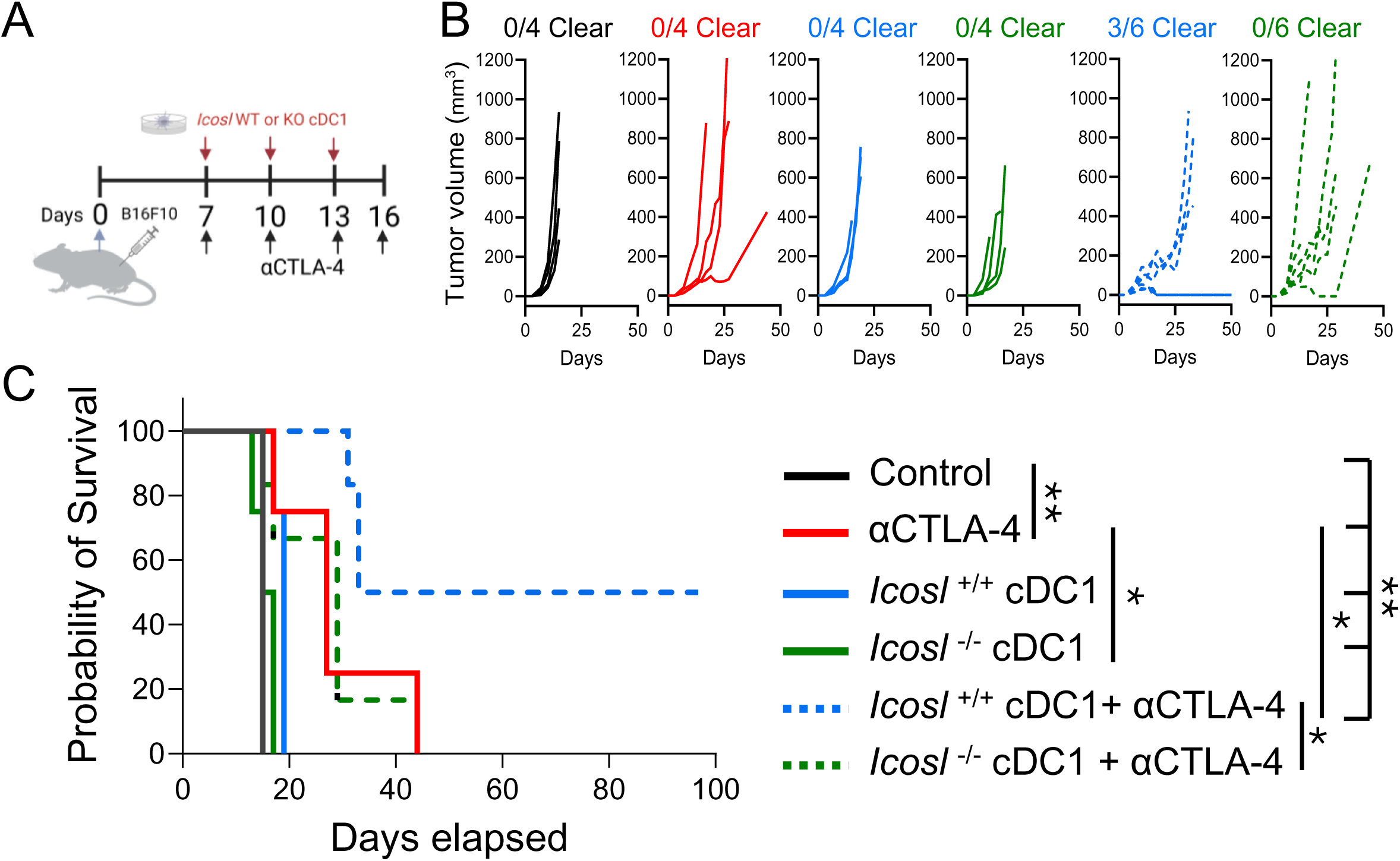
cDC1 ICOSL is required for anti-CTLA-4-mediated tumor eradication (related to figure 3) **(A-C)** *In vivo* adoptive transfer of cDC1 in combination with anti-CTLA-4 as outlined in **(A)**. Outgrowth of B16F10 tumors in B6 WT mice **(B)**. (n = 4-6 mice per group). Kaplan-Meier survival curves of tumor-bearing mice are shown **(C)**. Data are shown as means ± SEM. Log-rank (Mantel-Cox) test was used to determine statistical significance for survival of mice (**p<0.01, **p<0.01).

**Figure S5:**
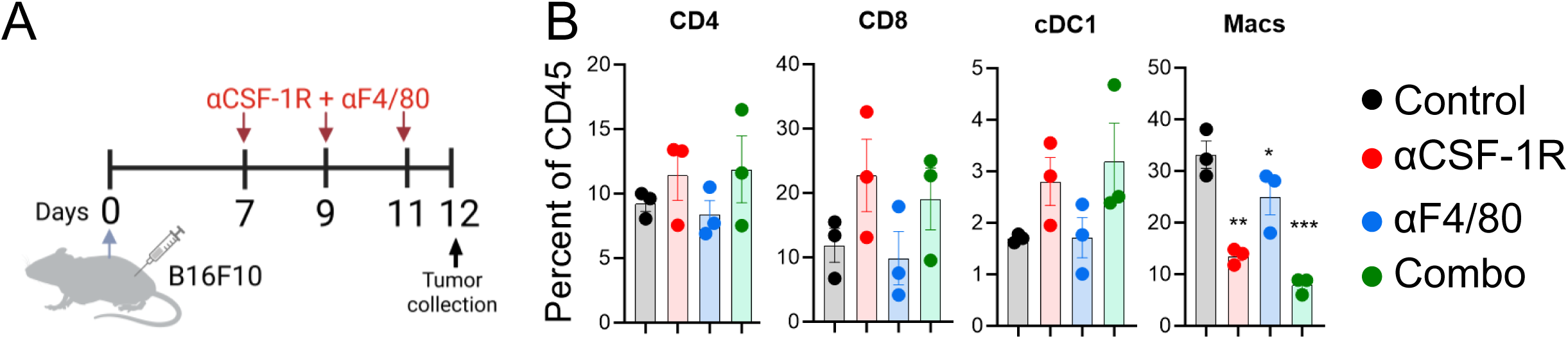
In vivo macrophage depletion (related to figure 4) **(A)** Experimental outline. **(B)** CD4^+^, CD8^+^ T-cell, cDC1 and macrophage frequencies in B16F10 tumors on day 12 as assessed by flow cytometry.

**Figure S6:**
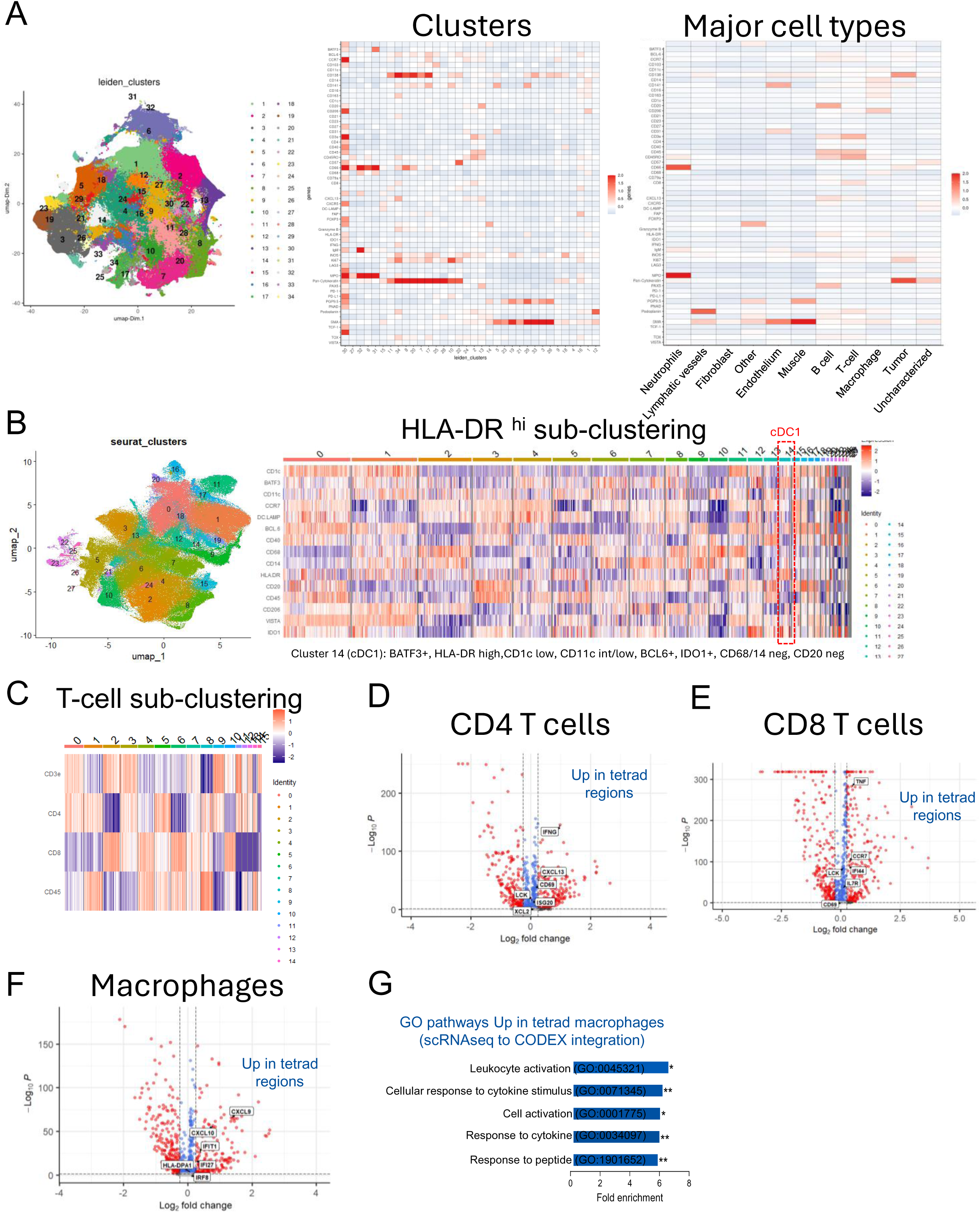
Processing and analysis of human CODEX data (related to figure 5) **(A)** Leiden clusters of major immune cell populations after pre-processing steps. **(B)** cDC1 sub-clustering from HLA-DR^hi^ cells. **(C)** CD4^+^ and CD8^+^ T-cell sub-clustering from CD3^+^ T-cells. **(D-F)** Volcano plot for DEGs between tetrad (red) and non-tetrad (blue) CD4^+^ T-cells (D), CD8^+^ T-cells **(E)** and macrophages **(F).** Statistically significantly DEGs (*p* < 0.05) are shown in red and blue, with select genes highlighted for reference. **(G)** Top 5 GO terms enriched for Visium HD genes upregulated in tetrad (red) macrophages.

## Notes

### Competing Interest Statement

The authors have declared no competing interest.

